# iTie controls intestinal tolerance through suppressing the NLRP6 inflammasome

**DOI:** 10.1101/2023.01.12.523719

**Authors:** Xiwen Qin, Fangrui Zhu, Weitao Li, Shuo Wang, Pengyan Xia

## Abstract

Intestines are full of commensal bacteria that possess numerous pathogen-associated molecular patterns. How the intestinal epithelial cells are tolerant to these stimuli under normal conditions is still elusive. Here we show that iTie is expressed in small intestinal enterocytes and its deficiency leads to body weight loss in mice, accompanied by length reduce of small intestines and intestinal villi. The activation of NLRP6 inflammasome is exacerbated upon *iTie* deletion. iTie has a higher binding affinity for NLRP6 than NLRP6’s physiological ligand LTA does. *iTie* deficiency gives rise to uncontrolled GSDMD activation and pyroptosis of small intestinal enterocytes. Inhibition of GSDMD permeabilization on cell membrane ameliorates the damage caused by *iTie* deficiency. iTe’s expression is diminished in small intestines of patients with Crohn’s disease. Our results uncover a self-control system for mouse intestine to tolerate commensal bacteria which might shed new light on the treatment of bowel diseases.

## Introduction

The intestinal microenvironment is filled with various commensal bacteria (Consortium, 2012; Martens et al., 2018; Sender et al., 2016). As the first barrier to contact with bacteria, intestinal epithelial cells play a key role in maintaining the stability of the intestinal mucosa (Allaire et al., 2018; Kayama et al., 2020). Among many mechanisms regulating intestinal epithelial homeostasis, the innate immune signaling pathway is of great significance (Perez-Lopez et al., 2016; Peterson and Artis, 2014). Too weak innate immune signals will cause the destruction of intestinal integrity and bacterial infection (Franchi et al., 2012; Knodler et al., 2014; Sellin et al., 2014; Sun et al., 2013). Too strong innate immune signals will also cause excessive inflammatory reaction, resulting in intestinal damage (Glocker et al., 2009; Guma et al., 2011; Lipinski et al., 2012; Xiao et al., 2007). Intestinal cells have developed a series of mechanisms consisting of physical barriers and appropriate cellular immune responses to tolerate intestinal commensal bacteria (Hooper et al., 2012; Patankar and Becker, 2020). However, how to balance the stimulation of commensal bacteria to intestinal cells and the strength of innate immune responses of intestinal cells is still not fully elucidated.

Inflammasome is a macromolecular protein complex, which usually contains a NOD-like receptor to sense danger signals from the outside or inside, an ASC protein that forms a scaffold and an executor caspase-1 (Christgen et al., 2020; Evavold and Kagan, 2019; Lamkanfi and Dixit, 2014; Rathinam and Fitzgerald, 2016). Activated caspase-1 can cut its substrate GSDMD (Broz et al., 2020; Wang et al., 2020). Matured GSDMD then form pores on the cell membrane, causing the release of inflammatory cytokines and pyroptosis (Ding et al., 2016; Liu et al., 2016; Shi et al., 2015). Components of the NLRP6 inflammasome are highly expressed in intestines and orchestrate the interface among host and microbes in the large intestine through controlling mucus secretion from goblet cells (Birchenough et al., 2016; Chen et al., 2011; Elinav et al., 2011; Levy et al., 2017; Wlodarska et al., 2014). Overactivation of the NLRP6 inflammasome arouses damaging colonic inflammation in mice challenged with pathogenic bacteria (Mukherjee et al., 2020). NLRP6 directly senses bacterium cell well component LTA or viral RNA (Hara et al., 2018; Shen et al., 2021; Wang et al., 2015). The function of NLRP6 in the large intestine has been widely reported, but NLRP6 is also highly expressed in the small intestine. The exact role of the NLRP6 inflammasome in small intestines remains elusive.

iTie, an intestinal tolerance implementer in enterocytes based on our study, is also known as NMES1, C15orf48 or AA467197 which is originally considered to be a nuclear protein negatively related to the occurrence of human esophageal oncogenesis (Zhou et al., 2002). iTie is also found to locate in the mitochondria, in where it replaces Ndufa4 during spermatogenesis of the reproduction system and inflammation of endothelial cells or primary macrophages (Clayton et al., 2021; Endou et al., 2020; Floyd et al., 2016; Lee et al., 2021). Compared with immune cells or endothelial cells, iTie is highly expressed in the esophageal system (Fagerberg et al., 2014). However, its role in intestinal cells remains unclear.

Here we show that iTie is critically involved in the maintenance of small intestinal tolerance. Mice deficient of *iTie* undergo body weight loss, accompanied by the shortening of small intestines and intestinal villi. Moreover, mice lacking *iTie* were more susceptible to infection by *L. monocytogenes* than wild type mice. The activation of NLRP6 inflammasome is enhanced in the absence of iTie. The binding affinity of iTie to NLRP6 was higher than that of LTA. *iTie* deficiency causes uncontrolled GSDMD activation and intestinal cell death. iTie suppresses NLRP6 inflammasome activation *in vitro* and *in vivo*. Inhibiting the permeability of GSDMD on cell membrane can save intestines from damages induced by *iTie* knockouts. In patients with Crohn’s disease, the expression of iTie in small intestines is reduced. Our findings reveal iTie as an implementer of the intestinal tolerance which may provide new clues for the treatment of intestinal diseases in the future.

## Results

### RNA sequencing identifies iTie as a potent implementer of the intestinal tolerance

We hypothesized that there were genes involved in the maintenance of intestinal tolerance to commensal bacteria. These genes should meet the following criteria: expressed in intestines, upregulated in the presence of commensal bacteria and downregulated in absence of microbes. To find these genes, we collected small intestines from postnatal mice that neither affected by the bacteria nor by the food, adult mice bred in germ-free or specific pathogen-free (SPF) conditions, and adult SPF mice treated with antibiotics. We then performed bulk RNA sequencing with these cells (Fig. 1A). There were eight genes that met the screening standards we set (Fig. 1B, C). We also validated the expression of these eight genes in small intestinal epithelium using the Single Cell Portal tools. Most of the eight genes were widely expressed throughout the intestine subsets while *AA467197* and *Tmem86b* were enriched in enterocytes (Fig. S1A). We then sought to find the physiological role of these genes *in vivo*. To generate gene knockouts in the intestine, we injected adeno-associated virus (AAV) particles that contain gRNAs specific for the eight genes to Rosa26-LSL-Cas9 knockin mice through the superior mesenteric artery from where AAV viruses are delivered to the intestinal epithelium (Fig. S1B). We monitored the body weight changes of these mice. We found that *AA467197* knockout induced the most significant body weight loss in mice (Fig. 1D). When we examined the digestive system one month later, we found *AA467197* deficiency significantly affected the length of small intestines (Fig. 1E), but did not affect the length of large intestines (Fig. S1C). Moreover, the length of small intestinal villi in *AA467197* knockout mice was also remarkably reduced (Fig. 1F and Fig. S1D), suggesting that *AA467197* is involved in intestinal homeostasis of mouse small intestines.

**Figure 1.**
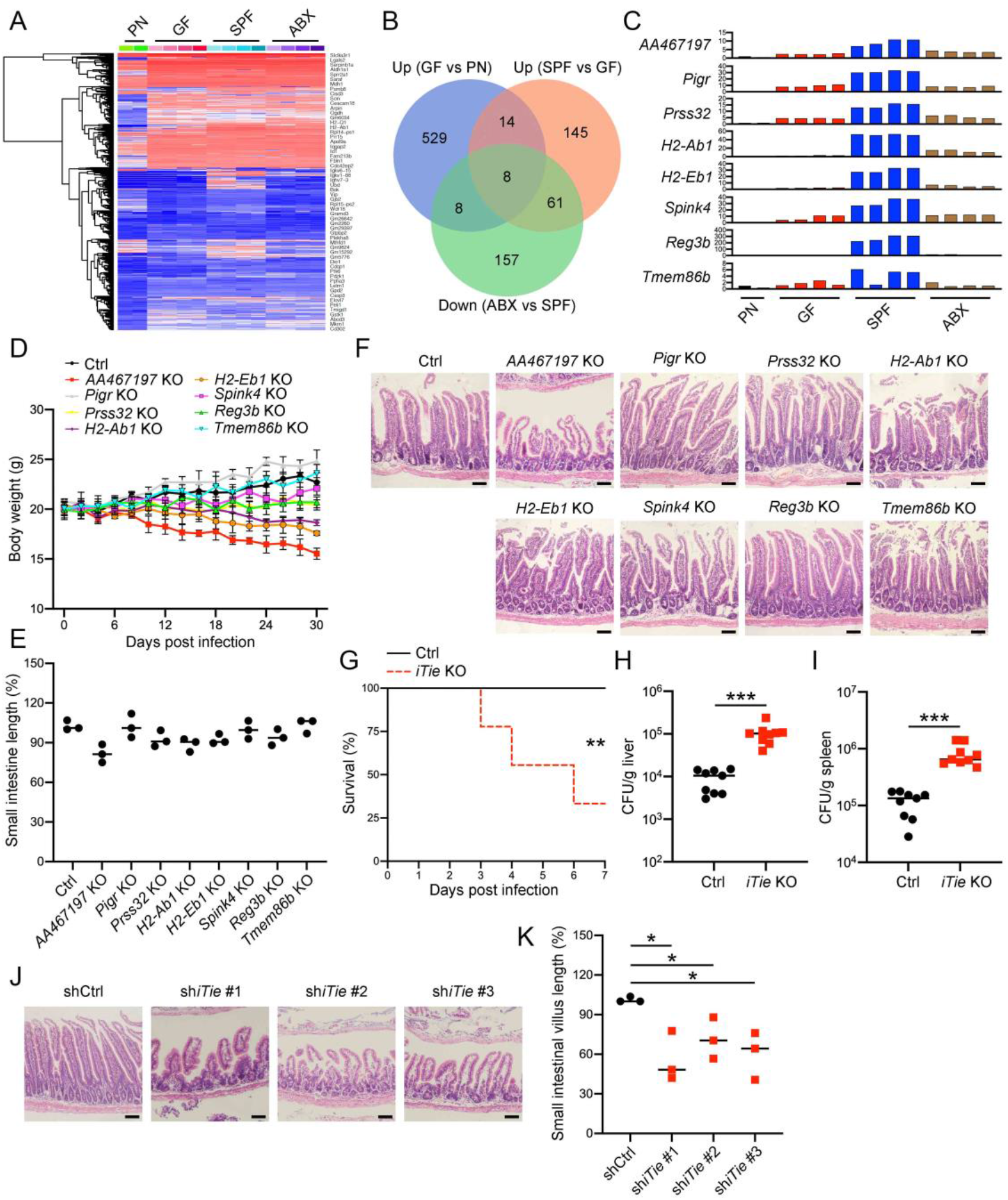
RNA sequencing identifies iTie as a potent implementer of the intestinal tolerance. (A) Small intestines from postnatal mice (PN), adult mice bred in germ-free (GF) or specific pathogen-free conditions (SPF), and adult SPF mice treated with antibiotics (ABX) were collected and subjected to bulk RNA sequencing. Expression levels of differential genes among the indicated groups were shown as heatmap. (B) A venn diagram was plotted using genes upregulated in the GF group compared with the PN one, genes upregulated in the SPF group compared with the GF one and genes downregulated in the ABX group compared with the SPF one. (C) Fold changes of expression levels of the indicated genes were calculated with the first sample of the PN group as 1. (D) Guide RNAs specific for the indicated genes together with a Cre recombinase were packaged into AAV (adeno-associated virus) viral particles. Viral particles were injected into Rosa26-LSL-Cas9 knockin mice through a superior mesenteric artery injection. Body weights were monitored in the following month. (E, F) One month post infection of gRNA-containing AAV viruses, mice were examined by quantifying the relative length of small intestines (E). Mouse small intestines were also sectioned and stained with hematoxylin-eosin (F). Scale bar, 50 μm. For D–F, three gRNAs were designed for each gene and one mouse was used for each gRNA. (G–I) One month post viral infections of control or *iTie* specific gRNAs, mice were intragastrically gavaged with 4 x 10^9^ *L. monocytogenes*. Survival of the indicated mice were calculated in the following 7 days (G, n=9). Bacterial loads in livers (H) and spleens (I) were examined 3 days post bacterial infection. AA467197 was renamed as iTie here. Three gRNAs were designed for *iTie* and three mice were used for each gRNA. (J, K) shRNAs specific for *iTie* were packaged into AAV viral particles, followed by viral particle delivery into mice through the superior mesenteric artery. Mouse small intestines were sectioned and stained with hematoxylin-eosin one month later (J) and the villus length changes were calculated between the control and *iTie* knockdown groups. Three shRNAs were designed for *iTie* and three mice were used for each shRNA. Scale bar, 50 μm. Data were shown as means±SD. *, *P*<0.05; **, *P*<0.01; ***, *P*<0.001. Experiments were repeated three times with similar results.

We thus focused on gene *AA467197* and renamed this gene as *iTie* (an Intestinal Tolerance Implementer in Enterocytes) based on its function we studied. To test whether *iTie* knockout had an effect on the host’s resistance to pathogenic bacteria, we challenged control and *iTie* knockout mice with *L. monocytogenes*. We found that lethal rates were increased in *iTie* KO mice when compared with control ones (Fig. 1G). Moreover, bacterial loads were increased in organs of *iTie* KO mice than that of controls (Fig. 1H, I). There is a microRNA (miR-147) that locates at the 3’ terminus of the *iTie* genetic locus. To exclude the possible impact of this microRNA on intestinal homeostasis, we generated AAV viruses containing shRNAs specific for the *iTie* mRNA. We then injected these viruses to wild-type mice and successfully knocked down *iTie* but not miR-147 (Fig. S1E, F). We found that mice with *iTie* knockdown had a body weight loss, length short of small intestines and intestinal villi, just as *iTie* KO mice did (Fig. S1G, H and Fig. 1J, K). Moreover, intragastrical infection of *L. monocytogenes* led to increased deaths in *iTie* knockdown mice, accompanied with bacterial loads raise (Fig. S1I–K). Altogether, these results suggest that iTie is a potent implementer of intestinal tolerance to microbes.

### *iTie* deficiency abrogates intestinal tolerance to commensal bacteria

To elucidate the physiological role of iTie in intestinal tolerance, we took use of a mouse strain that had the fourth exon of *iTie* deleted and a complete loss of the iTie protein (Fig. S2A). *iTie*^−/−^ mice showed significant weight loss (Fig. 2A). Moreover, the small intestine of *iTie*^−/−^ mice became shorter (Fig. 2B, C), while the length of the large intestine remained unchanged (Fig. 2D). Correspondingly, the small intestinal villi were shortened in *iTie* deficient intestines (Fig. 2E, F). We challenged *iTie*^+/+^ and *iTie*^−/−^ mice with *L. monocytogenes*. We found that lethal rates were elevated in *iTie*^−/−^ mice (Fig. 2G). Moreover, bacterial loads were increased in organs of *iTie*^−/−^ mice than that of *iTie*^+/+^ ones (Fig. 2H, I).

**Figure 2.**
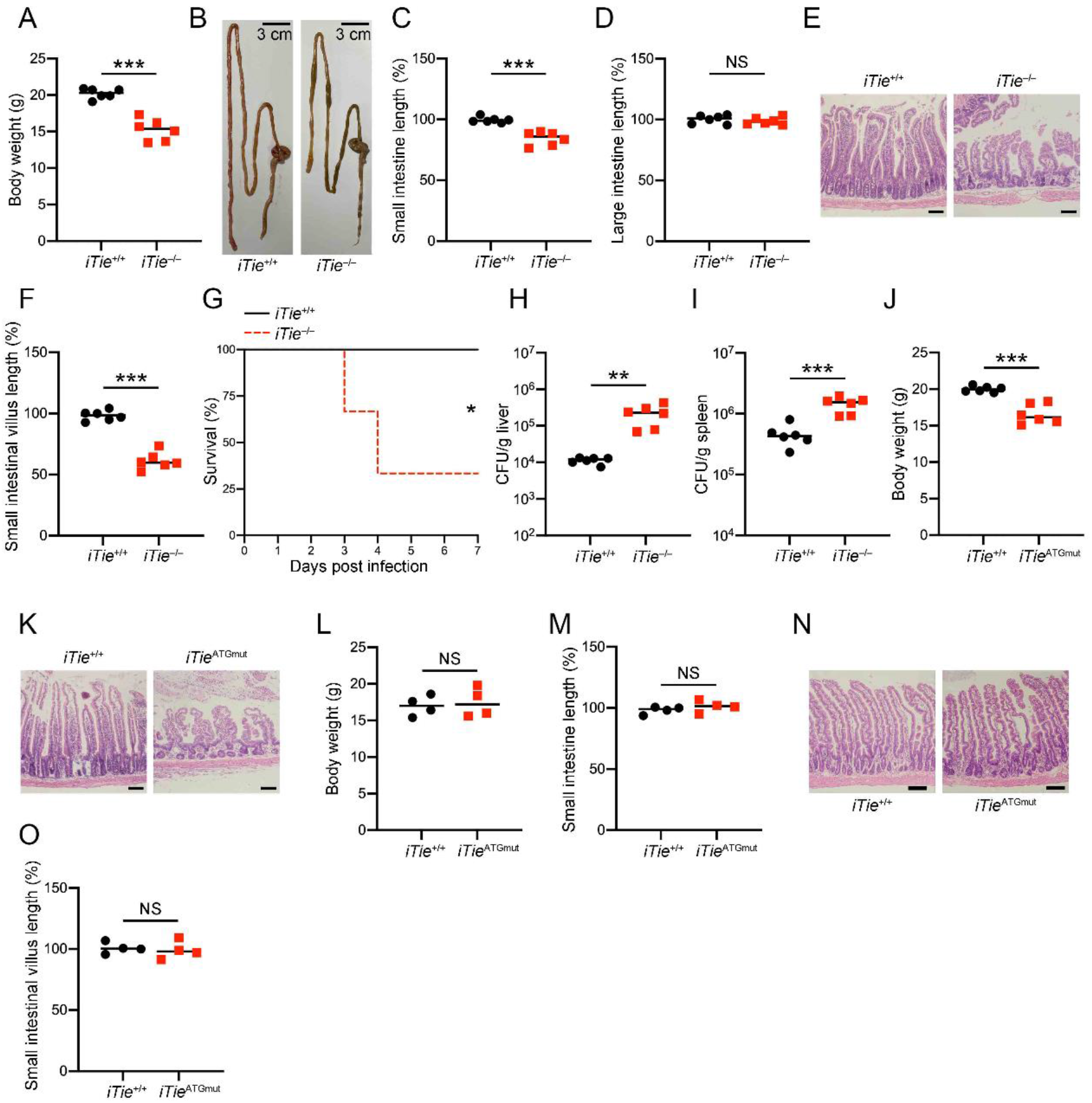
*iTie* deficiency abrogates intestinal tolerance to commensal bacteria. (A–D) Exon 4 of *iTie* was deleted through the CRISPR/Cas9 technology to generate *iTie*^−/−^ mice. *iTie*^+/+^ and *iTie*^−/−^ mice were bred and held in SPF conditions and body weights were checked at 8 weeks of age (A). Mouse guts were separated (B) and the relative length changes of small intestines (C) and large intestines (D) were examined. (E, F) Small intestines from *iTie*+/+ and *iTie*^−/−^ mice were sectioned and stained with hematoxylin-eosin (E). The relative length changes of small intestinal villi were calculated (F). Scale bar, 50 μm. (G–I) *iTie*^+/+^ and *iTie*^−/−^ mice were intragastrically gavaged with 4 x 10^9^ *L. monocytogenes*. Survival of the indicated mice were calculated in the following 7 days (G, n=6). Bacterial loads in livers (H) and spleens (I) were examined 3 days post bacterial infection. (J, K) The 8^th^ genetic codon of *iTie* was mutated to a stop codon (ATGmut) through the CRISPR/Cas9 technology. *iTie*^+/+^ and *iTie*^ATGmut^ mice were bred and held in SPF conditions and body weights were checked at 8 weeks of age (J). Small intestines from *iTie*^+/+^ and *iTie*^ATGmut^ mice were sectioned and stained with hematoxylin-eosin (K). Scale bar, 50 μm. (L–O) *iTie*^+/+^ and *iTie*^ATGmut^ mice were bred and held in SPF conditions with antibiotics and body weights were checked at 8 weeks of age (L). Small intestines of *iTie*^+/+^ and *iTie*^ATGmut^ mice were separated and the relative length changes were examined (M). Small intestines were also sectioned and stained with hematoxylin-eosin (N) and the relative length changes of small intestinal villi were calculated (O). Scale bar, 50 μm. NS, non-significant. Data were shown as means±SD. *, *P*<0.05; **, *P*<0.01; ***, *P*<0.001. Experiments were repeated three times with similar results.

To further exclude the influence of miR-147, we mutated the eighth genetic codon of the iTie ORF to a stop codon, leading to an early termination of the iTie peptides during translation (Fig. S2B, C). We did not mutate the translation start codon because mutation of the initial “ATG” during translation would cause an in-frame translation of the iTie ORF from the eighth genetic codon, which was also an “ATG”. These *iTie*^ATGmut^ mice had the same expression levels of *Mir147* as their wild-type counterparts (Fig. S2D). *iTie*^ATGmut^ mice showed a prominent body weight loss (Fig. 2J). The length of small intestine but not large intestine was diminished in *iTie*^ATGmut^ mice (Fig. S2E, F). Furthermore, the small intestinal villi were shortened in *iTie*^ATGmut^ mice (Fig. 2K and Fig. S2G). We then challenged *iTie*^+/+^ and *iTie*^ATGmut^ mice with *L. monocytogenes*. We found that *iTie*^ATGmut^ mice were more susceptible to bacterial infection in respect of decreased survival and increased bacterial loads in infected organs of *iTie*^ATGmut^ mice (Fig. S2H–J), suggesting that *iTie* deficiency hampers the integrity of the small intestine of mice infected with pathogenic microbes.

To validate that iTie also controls intestinal tolerance to commensal bacterial, we bred *iTie*^ATGmut^ mice in SPF conditions with antibiotics. Under such circumstances, *iTie*^ATGmut^ mice had the same body weight as their wild-type counterparts (Fig. 2L). Moreover, lengths of small intestines and large intestines were also comparable between *iTie*^ATGmut^ and control mice (Fig. 2M and Fig. S2K). The lengths of small intestinal villi were also unchanged post *iTie* deletion (Fig. 2N, O), suggesting that the alterations in small intestines of *iTie* KO mice bred in SPF conditions are induced by commensal bacteria.

We then co-bred newborn wild-type mice with *iTie*^ATGmut^ mice to check whether the phenomenon observed in *iTie*^ATGmut^ mice could be transferred to wild-type mice. Body weights were comparable between wild-type mice co-bred with *iTie*^ATGmut^ mice and ones co-bred with *iTie*^+/+^ mice (Fig. S2L). The lengths of small intestines and large intestines were also unchanged among these mice (Fig. S2M, N). Finally, small intestinal villi were not shortened in wild-type mice co-bred with *iTie*^ATGmut^ mice (Fig. S2O, P), suggesting that iTie sustains intestinal tolerance to commensal bacteria in a cell-intrinsic manner.

### iTie associates with NLRP6

To find how iTie participated in the maintenance of intestinal tolerance, we took use of the yeast two-hybrid system and screened a cDNA library derived from mouse enterocytes using iTie as a bait. 8 out of 9 positive clones screened out were identified as NLPR6 (Fig. 3A). We then verified the association of iTie and NLRP6 in HEK293T cells (Fig. 3B and Fig. S3A). We also generated recombinant iTie proteins fused with a GST tag. GST-iTie could pull down overexpressed NLRP6 in HEK293T cells (Fig. S3B). Moreover, GST-iTie but not GST itself could pull down endogenous NLRP6 from small intestinal lysates (Fig. 3C). Endogenous iTie was found to be associated with NLRP6 in enterocytes isolated from wild-type small intestines (Fig. 3D and Fig. S3C). We also mapped the binding regions between these two proteins. NLPR6 LRR domain was shown to be responsible for iTie binding and the iTie N terminus was essential for binding to NLRP6 (Fig. 3E, F). The interaction between iTie and NLRP6 was accelerated in enterocytes infected with *L. monocytogenes* (Fig. 3G and Fig. S3D). However, the association of iTie and NLRP6 was not affected by transfected poly(I:C) or LTA in HEK293T cells (Fig. S3E, F), suggesting that there might be other factors responsible for the increased binding of iTie with NLRP6 in enterocytes during infections.

**Figure 3.**
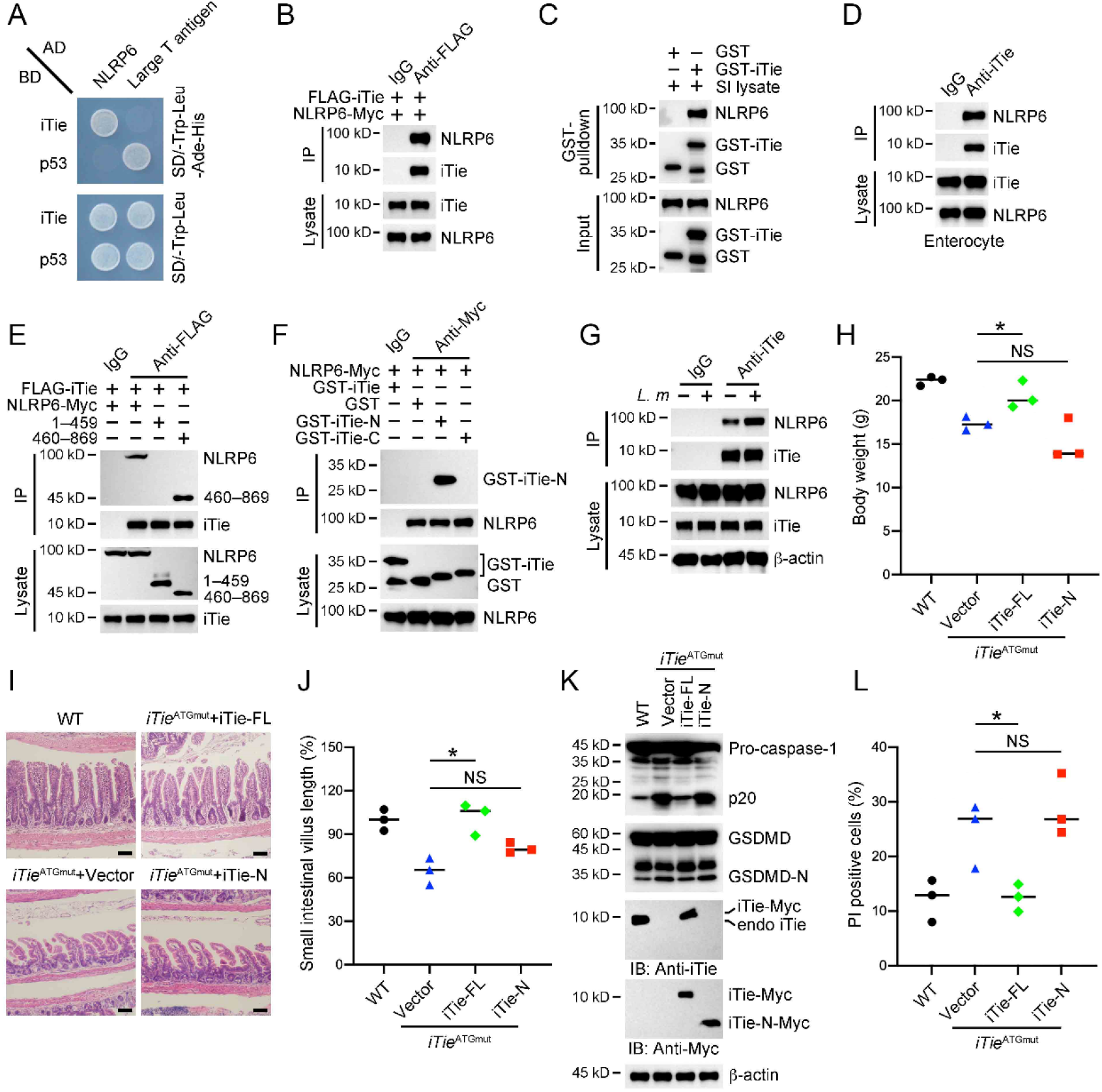
iTie associates with NLRP6. (A) Plasmids encoding Gal4 DNA binding domain (BD) tagged iTie and Gal4 activating domain (AD) tagged NLRP6 were co-transfected into yeast strain AH109. Transformants were grown in the indicated selection media. Clones transfected with p53 and large T antigen served as a positive control. (B) Plasmids encoding FLAG-tagged iTie and Myc-tagged NLRP6 were co-transfected into HEK293T cells for 24 h, followed by immunoprecipitation with antibody against FLAG or with a control IgG. Immunoprecipitations and cell lysates were immunoblotted with antibodies against the indicated proteins. (C) Recombinant GST-iTie or control GST protein were incubated with small intestinal lysates for 4 h, followed by a GST pull-down assay. Precipitates were immunoblotted with antibodies against the indicated proteins. (D) Enterocytes separated from the small intestine of WT mice were lysed and immunoprecipitated with antibody against iTie or with a control IgG. Precipitates were immunoblotted with antibodies against the indicated proteins. (E) Plasmids encoding FLAG-tagged iTie and Myc-tagged NLRP6 full-length or the indicated truncations were co-transfected into HEK293T cells for 24 h, followed by immunoprecipitation with antibody against FLAG or with a control IgG. Immunoprecipitations and cell lysates were immunoblotted with antibodies against the indicated proteins. (F) Recombinant GST-iTie or the indicated GST-fused truncations were incubated with HEK293T cell lysates containing Myc-tagged NLRP6 for 4 h, followed by immunoprecipitation with antibody against Myc or with a control IgG. Precipitates were immunoblotted with antibodies against the indicated proteins. (G) Enterocytes from the small intestine of WT mouse were incubated with *L. monocytogenes* at an MOI of 10 for 1.5 h. Cells were then supplemented with fresh medium containing 100 μg/ml Gentamycin and cultured for 16 h. Cells were then lysed and immunoprecipitated with antibody against iTie or with a control IgG. Precipitates were immunoblotted with antibodies against the indicated proteins. (H–J) AAV viral particles containing gene fragments of iTie full-length or iTie N terminus (aa 1–41) were delivered into *iTie*^ATGmut^ mice through the superior mesenteric artery. One month later, body weights were checked (H) and small intestines were sectioned and stained with hematoxylin-eosin (I). The relative length changes of small intestinal villi were also calculated (J). Scale bar, 50 μm. (K, L) Enterocytes were separated from the small intestines of iTie-FL or N terminus-rescued mice. Cells were either subjected to immunoblotting with antibodies against the indicated proteins (K) or stained with PI to calculate the percentages of PI positive cells (L). NS, non-significant. Data were shown as means±SD. *, *P*<0.05. Experiments were repeated three times with similar results.

We then rescued iTie and its N terminus in intestines of *iTie*^ATGmut^ mice using AAV viruses. Full-length iTie re-expression but not the N terminus regained the body weight in *iTie*^ATGmut^ mice (Fig. 3H), suggesting that iTie utilizes the full-length of its body to function normally. Moreover, the small intestine length was also restored in iTie rescued mice while the large intestine was not affected (Fig. S3G, H). Consistently, the length of small intestinal villi was elevated when iTie was reintroduced (Fig. 3I, J). We also checked the activation of NLRP6 inflammasome in these enterocytes restored with full-length iTie or its N terminus. We found that full-length iTie restoration suppressed the activation of caspase-1 and GSDMD when compared with cells deficient of iTie or rescued with iTie N terminus (Fig. 3K), suggesting that inflammasome overactivation in the absence of iTie might contribute to the disruption of intestinal tolerance to commensal bacteria. Moreover, the percentages of dying cells were also lowered in enterocytes possessing full-length iTie but not its truncations (Fig. 3L). Taken together, iTie associates with NLRP6 to modulate intestinal tolerance.

### *iTie* knockout leads to accelerated activation of the NLRP6 inflammasome

We then sought to decipher the role of NLRP6 inflammasome in iTie-mediated intestinal tolerance. Among the inflammasome related genes, components of the NLRP6 inflammasome were widely expressed in intestinal enterocytes (Fig. S4A). These genes were also constitutively expressed in the small intestines of postnatal, germ-free, SPF as well as antibiotics-treated mice (Fig. S4B). NLRP6 senses bacterial cell well component LTA. Indeed, inflammasomes were activated in enterocytes of SPF but not germ-free mice (Fig. S4C, D), indicating that the inflammasomes were persistently activated by commensal bacteria under SPF conditions. Moreover, *Nlrp6* deficiency abolished inflammasome activation and pyroptosis in enterocytes of mice bred in SPF conditions without antibiotics, further suggesting that NLRP6 is responsible for inflammasome activation in enterocytes (Fig. 4A, B). When we looked into the molecular status of *iTie* deficient cells, we found that there were intense inflammasome activation in these cells from mice bred without antibiotics only (Fig. 4C, D), implying that uncontrolled inflammasome activation might contribute to the shortened small intestines observed in *iTie* knockout mice and overactivation of the inflammasome destroys intestinal tolerance to commensal bacteria.

**Figure 4.**
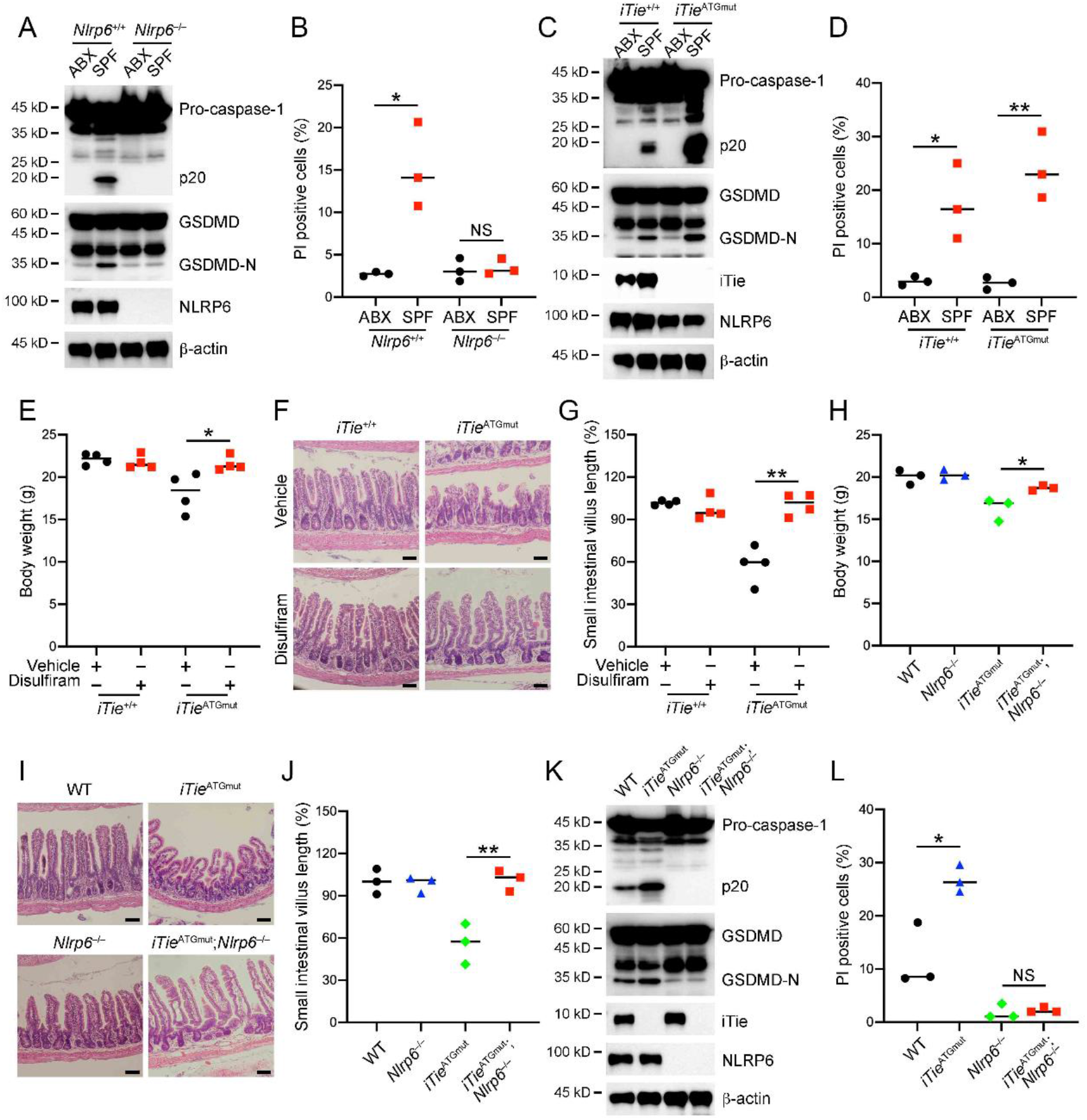
*iTie* knockout leads to accelerated activation of the NLRP6 inflammasome. (A, B) Enterocytes were isolated from the small intestines of *Nlrp6*^+/+^ and *Nlrp6*^−/−^ mice housed under SPF conditions with or without antibiotics. Cells were either subjected to immunoblotting with antibodies against the indicated proteins (A) or stained with PI to calculate the percentages of PI positive cells (B). (C, D) Enterocytes were separated from the small intestines of *iTie*^+/+^ and *iTie*^ATGmut^ mice bred under SPF conditions with or without antibiotics. Cells were either subjected to immunoblotting with antibodies against the indicated proteins (C) or stained with PI to calculate the percentages of PI positive cells (D). (E–G) *iTie*^+/+^ and *iTie*^ATGmut^ mice were administrated with 150 μg/ml disulfiram through drinking water for one month. Body weights were then checked (E) and small intestines were sectioned and stained with hematoxylin-eosin (F). The relative length changes of small intestinal villi were also calculated (G). Scale bar, 50 μm. (H–J) Body weights were determined in WT, *Nlrp6*^−/−^, *iTie*^ATGmut^ and *iTie*^ATGmut^; *Nlrp6*^−/−^ mice at 8 weeks of age (H). Small intestines from these mice were sectioned and stained with hematoxylin-eosin (I). The relative length changes of small intestinal villi were also calculated (J). Scale bar, 50 μm. (K, L) Enterocytes were separated from the small intestines of the indicated mice. Cells were either subjected to immunoblotting with antibodies against the indicated proteins (K) or stained with PI to calculate the percentages of PI positive cells (L). NS, nonsignificant. Data were shown as means±SD. *, *P*<0.05; **, *P*<0.01. Experiments were repeated three times with similar results.

Disulfiram was previously shown to inhibit pyroptosis through blocking GSDMD pore formation (Hu et al., 2020). We then gave disulfiram to *iTie* deficient mice through drinking water. Disulfiram administration ameliorated *iTie* deficient mice in respect of recovery of body weight and small intestine length (Fig. 4E and Fig. S4E), as well as growing small intestinal villi to normal lengths (Fig. 4F, G). Disulfiram treatment had little effect on large intestines (Fig. S4F). We also generated mice deficient for both *iTie* and *Nlrp6*. These double knockout mice had elevated body weight when compared with *iTie*^ATGmut^ mice (Fig. 4H). Small intestine lengths of double KO mice were also restored to normal levels while the large intestines were unaffected (Fig. S4G, H). Moreover, small intestinal villi grew to normal in mice deficient of *Nlrp6* (Fig. 4I, J). Consistently, inflammasome activation was abrogated in *Nlrp6* deficient enterocytes (Fig. 4K), accompanied with decreased dying cells (Fig. 4L). In all, our results suggest that *iTie* deficiency leads to accelerated activation of the NLRP6 inflammasome that is responsible for the disrupted intestinal tolerance to commensal bacteria.

### iTie renders diminished binding of NLRP6 to its ligands

We then reconstituted the NLRP6 inflammasome in HEK293T cells. We found that iTie significantly reduced NLRP6 inflammasome activation (Fig. 5A, B). iTie worked directly on NLRP6 for this suppression because iTie did not bind caspase-1 or GSDMD (Fig. S5A, B). There was an increasing level of inflammasome activation in *iTie* deficient cells and stimulation of enterocytes with NLRP6 ligand LTA or poly(I:C) further enhanced inflammasome activation (Fig. 5C and Fig. S5C), possibly explaining the reason that iTie deficient mice had a worse survival rate when mice encountered pathogenic microbes. To test whether iTie had a higher binding affinity for NLRP6 than LTA did, we incubated LTA with NLRP6 and iTie. LTA bound NLRP6 only in the absence of iTie (Fig. 5D), suggesting that iTie binds NLRP6 tighter than LTA does. We then measured the binding affinities between NLRP6 and iTie. We found that iTie indeed had a much higher binding affinity for NLRP6 than LTA did (Fig. 5E, F). What’s more, transfections of poly(I:C) or LTA into enterocytes did not alter the association between iTie and NLRP6 (Fig. S5D, E). We also examined how the NLRP6-ASC complex was influenced by iTie. We found that LTA accelerated the association between NLRP6 and ASC only in the absence of iTie (Fig. 5G) and ASC did not exist in the NLRP6-iTie complex (Fig. S5F). Finally, NLRP6 binding to LTA was increased in *iTie* deficient enterocytes (Fig. 5H) and ASC oligomerization was enhanced in enterocytes deficient of *iTie* (Fig. 5I). In sum, these results indicate that iTie directly suppresses the NLRP6 inflammasome.

**Figure 5.**
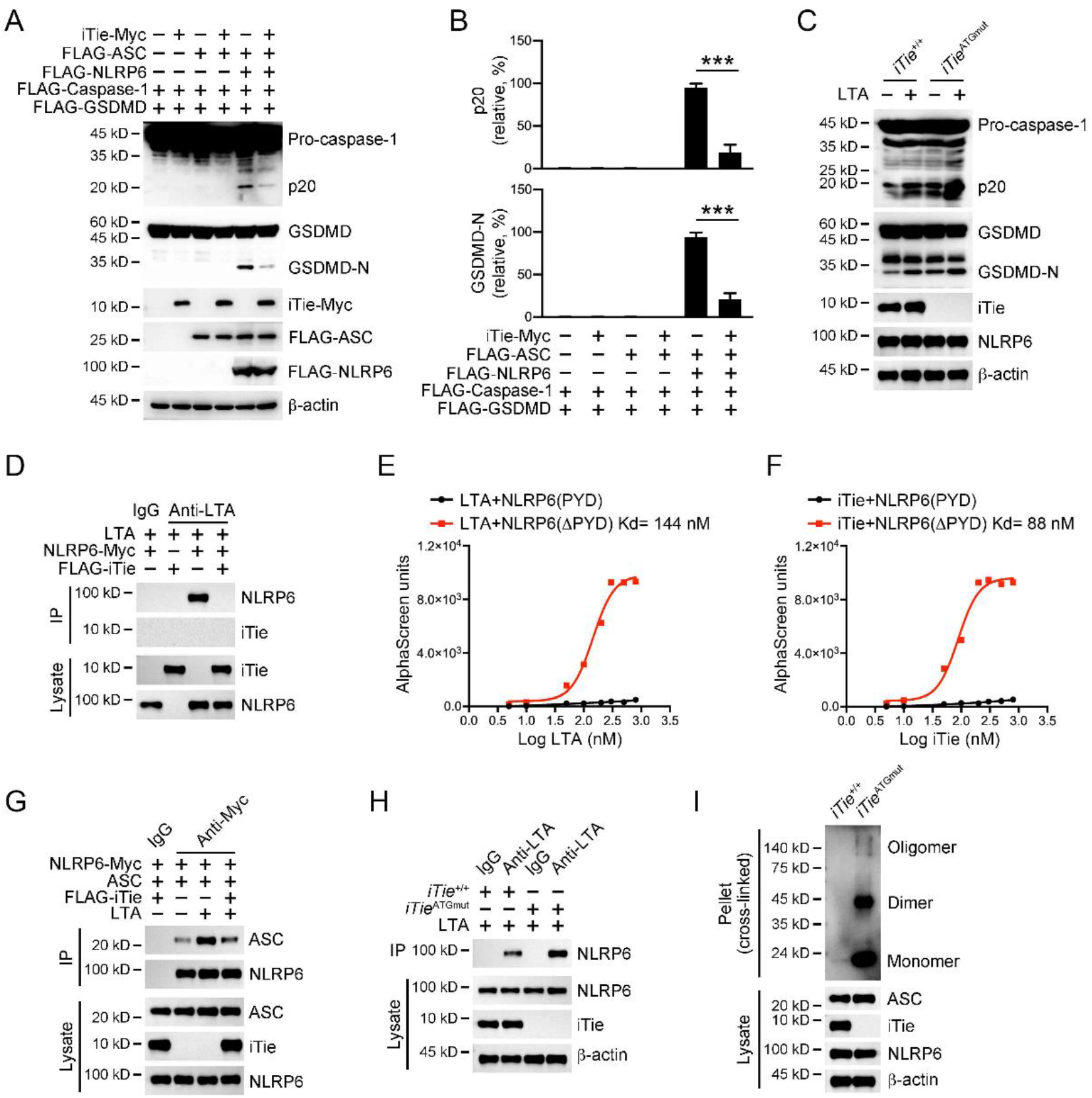
iTie hinders the binding of NLRP6 to its ligands. (A, B) Plasmids encoding Myc-tagged iTie, FLAG-tagged ASC, NLRP6, Caspase-1 and GSDMD were co-transfected into HEK293T cells for 24 h, followed by immunoblotting with antibodies against the indicated proteins (A). The cleaved band intensities of caspase-1 and GSDMD were quantified and shown as histograms (B). The fifth lane intensity was set as 1 and all others were compared to this one. (C) Enterocytes were separated from the small intestines of *iTie*^+/+^ and *iTie*^ATGmut^ mice. Cells were transfected with 10 μg/ml LTA for 4 h, followed by immunoblotting with antibodies against the indicated proteins. (D) Plasmids encoding FLAG-tagged iTie and Myc-tagged NLRP6 were co-transfected into HEK293T cells for 24 h. Cells were then lysed and incubated with 10 μg/ml LTA for 4 h, followed by immunoprecipitation with antibody against LTA or with a control IgG. Precipitates were immunoblotted with antibodies against the indicated proteins. (E) Plasmids encoding Myc-tagged NLRP6 (ΔPYD) or NLRP6 (PYD) were transfected into HEK293T cells for 24 h, followed by immunoprecipitation with magnetic beads coupled with antibody against Myc. Proteins were eluted by glycine. 400 nM Myc-tagged NLRP6 (ΔPYD) or NLRP6 (PYD) proteins were incubated with LTA of the indicated concentrations, followed by an AlphaScreen assay. (F) Plasmids encoding FLAG-tagged iTie, Myc-tagged NLRP6 (ΔPYD) or NLRP6 (PYD) were transfected into HEK293T cells for 24 h, followed by immunoprecipitation with magnetic beads coupled with antibody against FLAG or Myc. Proteins were eluted by glycine. 400 nM Myc-tagged NLRP6 (ΔPYD) or NLRP6 (PYD) proteins were incubated with iTie of the indicated concentrations, followed by an AlphaScreen assay. (G) Plasmids encoding FLAG-tagged iTie, Myc-tagged NLRP6 and ASC were co-transfected into HEK293T cells for 24 h. Cells were then transfected with 10 μg/ml LTA for 4 h, followed by immunoprecipitation with antibody against Myc or with a control IgG. Precipitates were immunoblotted with antibodies against the indicated proteins. (H) Enterocytes were separated from small intestines of *iTie*^+/+^ and *iTie*^ATGmut^ mice. Cells were transfected with 10 μg/ml LTA for 4 h, followed by immunoprecipitation with antibody against LTA or with a control IgG. Precipitates were immunoblotted with antibodies against the indicated proteins. (I) Enterocytes were separated from small intestines of *iTie*^+/+^ and *iTie*^ATGmut^ mice. Cells were then lysed and pellets were crosslinked with 2 mM DSS for 30 min at room temperature. Crosslinked pellets were dissolved in 1 x Laemmli buffer and immunoblotted with antibody against ASC. Cell lysates were also immunoblotted with antibodies against the indicated proteins. Data were shown as means±SD. ***, *P*<0.001. Experiments were repeated three times with similar results.

### *iTie* deletion exacerbates GSDMD-mediated pyroptosis of enterocytes

We then checked the activation status of GSDMD in *iTie* deficient mouse intestines. We found an enhanced GSDMD activation in *iTie* knockout intestines (Fig. 6A, B), accompanied by exacerbated pyroptosis (Fig. S6A). We also generated mice deficient for both *iTie* and *Gsdmd* and bred those mice under SPF conditions (Fig. S6B). *iTie* and *Gsdmd* double knockout mice had elevated body weight when compared with *iTie*^ATGmut^ mice (Fig. 6C). Small intestine lengths of those double KO mice were also restored to normal levels while the large intestines were hardly affected (Fig. S6C, D). Moreover, small intestinal villi grew to normal in mice deficient of *Gsdmd* (Fig. 6D, E). To test the role of commensal bacteria in GSDMD activation, we firstly bred *iTie* knockout mice in SPF conditions with antibiotics and then transferred them to SPF conditions without antibiotics. Contact of commensal bacteria led to accelerated GSDMD activation and cell pyroptosis in *iTie* deficient enterocytes (Fig. 6F, G). We then rescued GSDMD and its mutant in intestines of *iTie* and *Gsdmd* double knockout mice using AAV viral particles. Full-length GSDMD re-expression but not the mutant reduced the body weight in double knockout mice (Fig. 6H). Moreover, the small intestine length was also reduced in GSDMD rescued mice while the large intestine was not affected (Fig. S6F, G). Consistently, the length of small intestinal villi was diminished when GSDMD was reintroduced (Fig. 6I, J). In the last, we checked the inflammasome activation status in intestinal samples from patients with Crohn’s disease and healthy controls. iTie’s expression levels dropped in patients’ terminal ileum, along with enhanced inflammasome activation (Fig. 6K). Altogether, these results suggest that *iTie* deletion exacerbates GSDMD-mediated pyroptosis in enterocytes.

**Figure 6.**
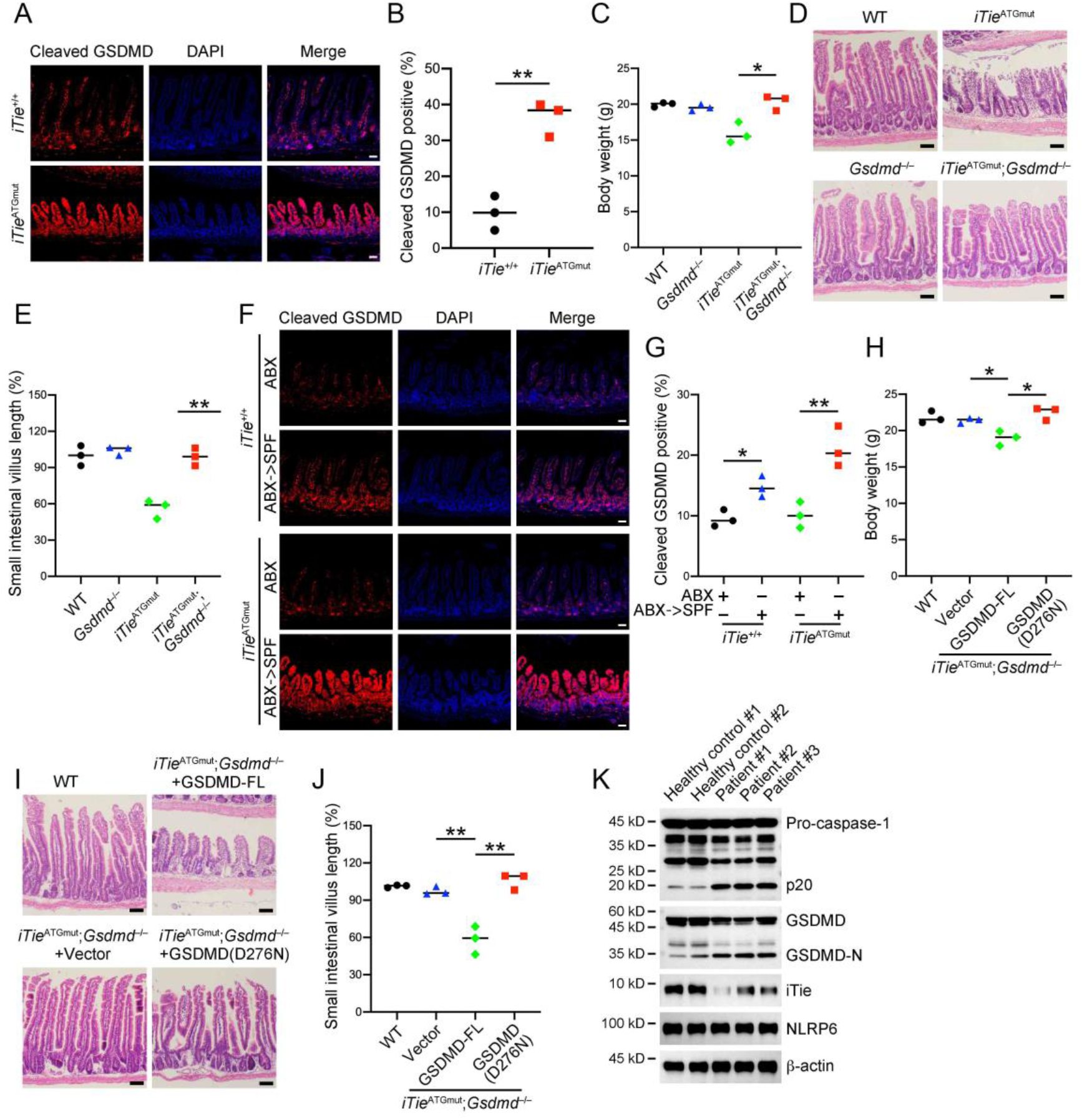
*iTie* deletion exacerbates GSDMD-mediated enterocyte pyroptosis. (A, B) Small intestines from *iTie*^+/+^ and *iTie*^ATGmut^ mice maintained in SPF conditions at 8 weeks of age were sectioned and stained with antibodies against cleaved GSDMD (A). The nuclei were counterstained with DAPI. Percentages of enterocytes with cleaved GSDMD signals were calculated (B). Scale bar, 50 μm. (C–E) Body weights were determined in WT, *Gsdmd*^−/−^, *iTie*^ATGmut^ and *iTie*^ATGmut^;*Gsdmd*^−/−^ mice at 8 weeks of age (C). Small intestines from these mice were sectioned and stained with hematoxylin-eosin (D). The relative length changes of small intestinal villi were also calculated (E). Scale bar, 50 μm. (F, G) 8 weeks-old *iTie*^+/+^ and *iTie*^ATGmut^ mice maintained in SPF conditions with antibiotics were transferred to SPF conditions without antibiotics for a month. Small intestines of the indicated mice were sectioned and stained with antibodies against cleaved GSDMD (F). The nuclei were counterstained with DAPI. Percentages of enterocytes with cleaved GSDMD signals were calculated (G). Scale bar, 50 μm. (H–J) AAV viral particles containing gene fragments of WT GSDMD or mutant GSDMD (D276N) were delivered into *iTie*^ATGmut^;*Gsdmd*^−/−^ mice through the superior mesenteric artery. One month later, body weights were checked (H) and small intestines were sectioned and stained with hematoxylin-eosin (I). The relative length changes of small intestinal villi were also calculated (J). Scale bar, 50 μm. (K) Enterocytes were separated from the terminal ileums of healthy controls or patients with Crohn’s disease, followed by immunoblotting with antibodies against the indicated proteins. NS, non-significant. Data were shown as means±SD. *, *P*<0.05; **, *P*<0.01; ***, *P*<0.001. Experiments were repeated three times with similar results.

## Discussion

Here we find an essential role of iTie in controlling intestinal tolerance to commensal bacteria. Without iTie, intestine enterocytes undergo elevated NLRP6 inflammasome activation when confronting microbes. iTie binds directly to the ligand binding domain of NLRP6 and constrains NLRP6 to avoid its overactivation in enterocytes. Thus, iTie contributes to the maintenance of intestinal tolerance to microbes.

Intestinal mucosae face various microbes and food allergens (Iweala and Nagler, 2019). Many mechanisms are reported to control intestinal tolerance to food allergens (Bryce, 2016; Okumura and Takeda, 2017; Peterson and Artis, 2014). Unlike food allergens that are absorbed by intestines, commensal bacteria in the intestine are rarely found inside the inner side of intestines under normal conditions (Perez-Lopez *et al*., 2016). In the large intestine, mucins secreted by the goblet cells provide a barrier to separate the commensal microbiota from intestinal cells (Johansson et al., 2011; Martens *et al*., 2018). There are also numbers of genes involved in intestinal tolerance maintenance and deficiency of which usually leads to intestinal over-reactivities to microbes either commensal or pathogenic (Mowat, 2018). In this study, we show that the small intestinal tolerance is maintained through suppressing the innate immune response in enterocytes. iTie is expressed in intestinal enterocytes and its deficiency leads to the disruption of intestinal tolerance to commensal bacteria. *iTie* deficient intestines have normal morphology in mice bred under conditions treated with antibiotics, further supporting our finding that iTie is required for the maintenance of intestinal tolerance in the intestines. Besides commensal bacterial, *iTie* knockout mice are also susceptible to pathogenic microbe infections, hinting that an undermined tolerance in these *iTie* deficient enterocytes.

NLRP6 has been shown to regulate mucin secretion in goblet cells of large intestines (Birchenough *et al*., 2016; Wlodarska *et al*., 2014). The NLRP6 inflammasome signaling also participates in shaping intestinal microenvironments when microbiota-modulated metabolites are present (Levy et al., 2015; Li and Zhu, 2020). Overactivation of the NLRP6 inflammasome leads to colonic inflammation under pathogenic conditions (Mukherjee *et al*., 2020). Here we find that iTie is co-expressed with components of the NLRP6 inflammasome in small intestinal enterocytes. iTie binds to NLRP6’s ligand binding domain and expels microbial ligands from NLRP6. iTie constrains the activation of NLRP6 inflammasome under normal states and maintains intestinal tolerance to commensal bacteria. *iTie* deficiency specifically affects the small intestine but not the large intestine. One possible explanation is that accelerated NLRP6 inflammasome activation in *iTie* knockout mice makes goblet cells secret more mucins that separate microbes from large intestinal enterocytes to quench NLRP6 inflammasome activation. The detailed mechanism underlying this needs to be further investigated.

In sum, we find an essential role of iTie in controlling intestinal tolerance through suppressing the NLRP6 inflammasome. Our finding will shed new light on the treatment of intestinal diseases such as Crohn’s disease in the future.

## Experimental procedures

### Antibodies and reagents

Antibodies against caspase-1 (3866, 24232 and 2225), ASC (13833, 67824) were from Cell Signaling Technology. Antibody against caspase-1(ab207802) was purchased from Abcam. Antibodies against NLRP6 (PA521022), lipoteichoic acid (MA1-40134) and iTie (PA5-69665) were from Invitrogen. Antibodies against NLRP6 (SAB1302240) and iTie (HPA012943) were from Sigma-Aldrich. Antibodies against lipoteichoic acid (NBP1-60146) and iTie (NBP1-98391) were from Novus Biologicals. Rabbit antibody against mouse iTie was generated by SinoBiological, Beijing. Anti-c-Myc magnetic beads (88843) were purchased from Pierce. Antibody against 6xHis tag (MA121315) was purchased from Invitrogen. Antibodies against FLAG tag (F3165), β-actin (A1978) and anti-FLAG M2 magnetic beads (M8823) were purchased from Sigma-Aldrich. Antibodies against c-Myc (sc-40) and GST (sc-138) were purchased from Santa Cruz Biotechnology. Anti-HA tag antibody (HX1820) was purchased from Huaxingbio (Beijing). HRP-conjugated goat antimouse IgG (SA00001-1) and HRP-conjugated goat anti-rabbit IgG (SA00001-2) were purchased from Proteintech Group. Alexa Fluor Plus 488-conjugated goat anti-mouse IgG (A32723) and Alexa Fluor 594-conjugated goat anti-rabbit IgG (A32740) were purchased from Invitrogen. Alexa Fluor Plus 488-conjugated donkey anti-goat IgG (bs-0294D-AF488) and Alexa Fluor 594-conjugated donkey anti-rabbit IgG (bs-0295D-AF594) were purchased from Bioss (Beijing).

Glutathione sepharose 4B resin (17075601) was purchased from Cytiva. Protein A/G PLUS-Agarose beads (sc-2003) were purchased from Santa Cruz Biotechnology. DAPI (2879038) was purchased from PeproTech (BioGems). Dropout (DO)/–Leu/–Trp and DO/–Ade/–His/–Leu/–Trp supplement, SMART MMLV reverse transcriptase (639524) and carrier DNA (630440) were purchased from Clontech, Takara Bio. Random primer (Hexadeoxyribonucleotide mixture, pd(N)6) (3801) was purchased from Takara Bio. Protease inhibitor cocktail (11697498001) was purchased from Roche. Total RNA extraction kit (8034111) was purchased from Dakewe (Beijing). Ni-NTA agarose beads (R90115) and AminoLink coupling resin (20381) were purchased from Invitrogen. DOTAPs were: D10530 from Psaitong (Beijing) and 11202375001 from Roche. Poly(I:C) (low molecular weight, tlrl-picw) and LTA (tlrl-pslta) were purchased from InvivoGen. Disulfiram (PHR1690) propidium iodide (P4170) were purchased from Sigma-Aldrich.

### Mice and infections

*Nlrp6*^−/−^ (S-KO-00213), *Gsdmd*^−/−^ (S-KO-12963) and *iTie^−/−^* (S-KO-10082) mice were purchased from Cyagen Biosciences (Jiangsu, China). Rosa26-LSL-Cas9 knockin mice (024857) were purchased from Jackson Laboratory. *iTie*^ATGmut^ mice were generated by Cyagen Biosciences. Briefly, the 8^th^ genetic codon (ATG) of *iTie* was mutated to a stop codon (TAG) through the CRISPR/Cas9 technology. Guide RNAs targeting exon 2 of mouse *iTie* gene, donor oligo with targeting sequence flanked by about 60 bp homologous sequences on each end, and Cas9 mRNA were introduced into fertilized mouse eggs through microinjection to generate *iTie* mutant offsprings. F0 founder mice were identified through sequencing and crossed with wildtype mice to generate F1 generation. Guide RNA sequences were: gRNA1, 5’-CCTTATTCTTCATCAATATC-3’; gRNA2, 5’-CCAGATATTGATGAAGAATA-3’. Mutant mice were identified through sequencing. Donor oligo was: TTTTCCTTCTTCTAGATTTGGAATTCTACACTAAAGTCATCATGGGCGTTTTCCAGATATTGTAGAAGAATAAGGAAGTAAGTTTCTAATCTTAACTTCTGAGAGTTTTCCTTTTTAAAAA. Mice were bred and held in specific pathogen-free conditions if not specified. Germ-free mice were purchased from Cyagen Biosciences and maintained at the department of laboratory animal science of Peking University. For survival calculations, mice were intragastrically gavaged with 4 x 10^9^ *L. monocytogenes* (strain 10403s). Mice survival were calculated in the following 7 days. Bacterial loads in livers and spleens were examined 3 days post bacterial infection.

### Primary cell separation and culture

For enterocytes separation, mouse small intestines were isolated and washed in icecold PBS three times, followed by slicing into small pieces about 1–2 mm in length. Tissue fragments were placed into ice-cold PBS containing 20 mM EDTA and shook for 1 h. Tissues were then shaken violently and supernatants were collected. Cells were stained with fluorescent-conjugated antibodies against CD45, CD31, TER119 and EpCAM. CD45^−^CD31^−^TER119^−^CpCAM^+^ cells were collected. Cells were cultured in mixed media composed of DMEM and F12 in a 1:1 ratio. Media were further supplemented with 5% FBS and 1% L-glutamine. Cells were cultured at 37 °C with a 5% CO_2_ humidified atmosphere. Small intestine samples from patients with or without enteritis were rinsed in ice-cold PBS and lysed for immunoblotting. Informed consents were obtained from all subjects and experiments conformed to related principles. Study was licensed by the Ethics Committee of Institute of Microbiology, Chinese Academy of Sciences.

### Histology and villus length calculation

Intestines were opened longitudinally and fixed in 4% PFA (paraformaldehyde, Sigma) for 10 h, followed by rehydration in 30% sucrose for 36 h. Tissues were immersed into paraffins and sectioned, followed by Hematoxylin and Eosin staining. The lengths of small intestine villi at the same position of the intestinal tract between experimental groups and controls were measured.

### Cell transfection

Transfection reagent JetPRIME (Polyplus Transfection) was used to conduct transfection in HEK293T cells. HEK293T cells were plated 14 h earlier to achieve a density of about 70% when transfection was performed. DOTAP was used for transfecting LTA or poly(I:C) into HEK293T or intestinal enterocytes.

### AAV virus packaging and purification

Guide RNAs or shRNAs under a U6 promoter and exogenous genes under a CMV promoter were cloned into AAV-vectors. AAV-vector, pANC80L65 and AAV-helper were transfected into HEK293T cells at a ratio of 1:1:2. 48 h later, supernatants were collected and precipitated with PEG8000. At the meantime, cell pellets were sonicated and mixed with the precipitates from the cell culture supernatants. Mixtures were centrifuged in a gradient iodixanol solution at 48,000 rpm for 2.5 h at 37 °C. Fractions between the 40% and 60% solution were collected and diluted in PBS buffer containing 0.001% pluronic F-68. Iodixanol was removed through buffer exchange in a 100 kD Amicon ultra centrifugal filter unit (Merck Millipore). Titers of AAV viruses were determined through transducing HEK293T cells of diluted viruses.

### Intestinal knockout and rescue

An AAV vector AAV:ITR-U6-sgRNA-hSyn-Cre expressing CRISPR-Cas9 targeting guide RNA (gRNA) designed for target genes and a Cre recombinase were used. Guide RNAs were designed using the Sequence scan for CRISPR of Harvard University. gRNAs were as follows: 5’-GAAAGATGTGGCTCCGGTGG-3’, 5’-AGCGAAAGATGTGGCTCCGG-3’ and 5’-CCTTATTCTTCATCAATATC-3’ for *AA467197*; 5’-TGGTGCCGACAAGGAGCCAG-3’, 5’-CCGGGTGTGCCGGTTGACAG-3’ and 5’- GGCCTGGGTACCAGTAACCG-3’ for *Pigr*; 5’-CGTCTACCACGACAGGCAGG-3’, 5’-AAGATGCCCAGCTGGGCCGG-3’ and 5’-AATGGGGCGCACGTGTGTGG-3’ for *Prss32*; 5’-CAGATACATCTACAACCGGG-3’, 5’-CGACGTGGGCGAGCACCGCG-3’ and 5’-GGAGATCCTGGAGCGAACGC-3’ for *H2-Ab1*; 5’-CGACGTGGGCGAGTTCCGCG-3’, 5’-TCCGCGCGGTGACCGAGCTG-3’ and 5’-GCGCGGTGACCGAGCTGGGG-3’ for *H2-Eb1*; 5’-AAGGAGGGGAACCAAGGTCA-3’, 5’-CAGACACCCAACCTGATCTG-3’ and 5’-ACCCAACCTGATCTGTGGCA-3’ for *Spink4*; 5’-CATCCAGGACATGACGGAGC-3’, 5’-GGGGCAACTAATGCGTGCGG-3’ and 5’-GCAGTAGGAGCCATAAGCCT-3’ for *Reg3b*; 5’-GGTTGTGTTCCTGTGGGCTG-3’, 5’-TGAAGGGGGCCAGCCACCTG-3’ and 5’-GGAGCAGGCAAGGATGAAGG-3’ for *Tmem86b*; 5’-GGGCGAGGAGCTGTTCACCG-3’, 5’-CGAGGAGCTGTTCACCGGGG-3’ and 5’-GAAGGGCATCGACTTCAAGG-3’ for eGFP. shRNAs were as follows: 5’-TTATTCTTCATCAATATCC-3’, 5’- TATACAAAGCGAAAGATGC-3’, 5’-TTTATTCTTCATCAATATC-3’ for iTie; 5’-TTGATATAGACGTTGTGGC-3’, 5’-TATGATATAGACGTTGTGC-3’, 5’-TTGAAGTTCACCTTGATGC-3’ for eGFP. SMA injection was performed as described previously (Polyak et al., 2008). 1 x 10^11^ AAV particles in 100 μl PBS were delivered into Rosa26-LSL-Cas9 knockin mice, or into mice deficient of *iTie* or *Gsdmd*.

### Inflammasome induction and reconstitution

Enterocytes isolated from mouse small intestines were incubated with *L. monocytogenes* at an MOI of 10 for 1.5 h. Cells were then supplemented with fresh medium containing 100 μg/ml Gentamycin and cultured for 16 h. For NLRP6 inflammasome reconstitution *in vitro*, plasmids encoding Myc-tagged iTie, FLAG-tagged ASC, NLPR6, Caspase-1 and GSDMD were co-transfected into HEK293T cells for 24 h, followed by examination of caspase-1 and GSDMD through immunoblotting.

### RNA extraction and RT-PCR

Total RNA extraction kit (Dakewe, Beijing) was used to extract RNAs in tissue cells or cultured cells according to manufacturer’s instructions. For reverse transcription, 1 μg RNA was mixed with 2 μl N6 primers and 7 μl DEPC water, followed by incubation at 72 °C for 2 min. The mixture was then chilled on ice for 2 min and then mixed with 4 μl 5x first-strand buffer (Takara Bio), 2 μl 100 mM DTT, 2 μl 10 mM dNTPs mix (Solarbio, Beijing) and 2 μl SMART MMLV reverse transcriptase (Clontech, Takara Bio). The mixture was incubated under the following conditions: at 25 °C for 10 min, at 42 °C for 1 h and at 75 °C for 10 min. cDNA samples were diluted for further PCR analysis. Primers for PCR identification were: 5’-ATTTGGAATTCTACACTAAAGTC-3’ (forward) and 5’- ACCACGTCGGTTTTTTTCAAC-3’ (reverse) for mouse *iTie*; 5’-TATGAATCTAGTGGAAACATTTC-3’ (forward) and 5’- CCTACAAATGTAGCAGAAGCA - 3’ (reverse) for mouse *Mir147*; 5’-ACACTCTCCCAGAAGGAGGG-3’ (forward) and 5’- TTTATAGGACGCCACAGCGG-3’ (reverse) for mouse *Actb*.

### Recombinant protein expression and purification

For recombinant expression of GST-tagged iTie, iTie cDNA was cloned into pGEX-6P-1 vector. Plasmids were transformed into *E. coli* strain BL21 and the bacteria grew in LB medium supplemented with 100 μg/ml ampicillin at 37 °C for 8 h, followed by addition of IPTG to a final concentration of 0.5 mM and shaking at 150 rpm at 37 °C for 3 h. Cells were harvested and resuspended with PBS containing 1% Triton-X-100 and lysed by sonication. Cell lysates were centrifuged at 13,000 rpm at 4 °C for 30 min and filtered through sterile syringe filters. The filtered lysate was loaded on column containing glutathione sepharose 4B resin, followed by washing with PBS containing 1% Triton-X-100. The protein was eventually eluted by 1 ml 10 mM L-Glutathione solution.

### Immunoprecipitation

HEK293T cells were seeded on 6-well culture plates and were transfected with plasmids 16 h later. 24 h post transfection, cells were washed twice with PBS and lysed in a pre-chilled buffer containing 150 mM NaCl, 50 mM Tris (pH 7.5), 1 mM EDTA, 0.5% digitonin, 1% protease inhibitor cocktail and 10% glycerol on ice for 30 min. For immunoprecipitation using primary cells, cells post stimulation or treatments were lysed in a pre-chilled PBS buffer containing 0.5% digitonin, 1% protease inhibitor cocktail on ice for 30 min. Cell lysates were centrifuged at 13,000 rpm at 4 °C for 10 min. Primary antibodies and their isotype control IgG were immobilized to AminoLink Coupling Resin (ThermoFisher Scientific) following the manufacturer’s instructions. Supernatants were incubated with immobilized antibodies at 4 °C for 2 h. Resins were washed three times with PBS containing 0.5% digitonin, followed by immunoprecipitant detachment in 0.1 M Glycine-HCl, pH 2.7. Supernatants were neutralized by adding 1/10 the volume of 2 M Tris-HCl, pH 8.0, followed by immunoblotting with the indicated antibodies.

### Immunofluorescence

After antigen retrieval, intestine sections were washed twice with PBS and blocked in 10% normal goat serum at 37 °C for 30 min. Samples were incubated with primary antibody for 2 h, followed by washing with PBS for three times and further incubation with fluorescence-conjugated secondary antibody for 1 h. Nuclei were stained with DAPI. Cells were visualized through an UltraVIEW VoX imaging system (PerkinElmer).

### RNA sequencing and analysis

For ABX treatments, SPF wild-type mice at 8 weeks of age were administrated with drinking water containing 1 mg/ml ampicillin, 1 mg/ml gentamicin, 1 mg/ml metronidazole, 1 mg/ml neomycin, 0.5 mg/ml vancomycin, 5 g/L sucrose for one month. Small intestines of newborn mice, germ-free and SPF mice at 8 weeks of age and ABX mice were collected. Tissues were homogenized and resuspended in TRIzol for total RNA extraction. mRNA was purified from total RNA through poly-T oligo-conjugated magnetic beads. Fragmentation buffer was added to break the mRNA into short fragments. First strand cDNA was synthesized through random hexamer primers and RNase H. Second strand cDNA was synthesized later. Double strand cDNA was purified by AMPure P beads and repaired by adding a tail at the end. Products were amplified by PCR. Clustering of the index-coded samples were performed on a cBot cluster generation system using HiSeq PE Cluster Kit v4-cBot-HS (Illumina) according to the manufacturer’s instructions, followed by sequencing on an illumine novaseq 6000 S4 platform and 2 x 150 bp paired-end reads were generated. Sequenced reads were aligned to mouse genomic reference GRCm38.84 using HISAT2 v2.1.0, followed by sorting using samtools v1.10. Sorted BAM files were then processed by StringTie v2.1.1 and normalized expression values were obtained. Differential genes among groups were selected using the following criteria: expression value above 1 and fold changes above 2. Genes upregulated in germ-free mice versus postnatal mice, genes upregulated in SPF mice versus germ-free mice and genes downregulated in ABX mice versus SPF mice were used to plot heatmaps and venn diagram. Sequencing data and processed expression matrix had been deposited into EBI ArrayExpress with an accession number E-MTAB-12215.

### Yeast two-hybrid screening

Yeast strain AH109 was inoculated into 5 ml YPDA medium and shaked at 250 rpm at 30 °C for 16 h. The overnight culture was transferred to a flask containing 50 ml YPDA and incubated at 30 °C for 3 h. Yeast cells were collected by centrifuged at 4,000 rpm for 10 min and resuspended in a 1XTE/LiAc buffer (pH7.5) containing 0.1 M lithium acetate, 0.1 M Tris-HCl and 10 mM EDTA. A mixture of plasmid DNA and carrier DNA was added to the yeast cell suspension and vortexed to mix well. A PEG/LiAc buffer containing 40% polyethylene glycol 3350, 0.1 M lithium acetate, 0.1 M Tris-HCl and 10 mM EDTA was then added to the suspension and vortexed at a high speed. The suspension was shaked at 200 rpm at 30 °C for 30 min, followed by addition of DMSO and heat shock in a 42 °C water bath for 15 min. Cells were chilled on ice for 2 min and resuspended to plated on appropriate SD agar plates. A mouse enterocyte cDNA library was used for Y2H screening with Gal4 DNA binding domain (BD)-tagged iTie as a bait. BD-tagged f iTie and Gal4 DNA activating domain (AD)-tagged NLRP6 were co-transfected into yeast strain AH109 and double positive clones were selected on SD medium lacking adenine, histidine, leucine and tryptophan.

### Statistical analysis

No statistical methods were used to predetermine sample size. Experiments were independently repeated at least three times to achieve statistical significance. No randomization or blinding procedures was used in this study. No samples were excluded from the analysis. Data were shown as means ± SD of three technical replicates. Data with normal distribution determined by Shapiro–Wilk normality test were statistically analyzed by two tailed Student’s t tests if not specified. The Gehan-Breslow-Wilcoxon test was used for the analysis of survival data. Data were analyzed by GraphPad Prism 9.0. *P*-values ≤0.05 were termed as significant (*, *P*<0.05; **, *P*<0.01; ***, *P*<0.001); NS means non-significant where *P*>0.05.

## Acknowledgements

We thank Ting Li (Peking University) for technical help. This work was supported by the National Key R&D Program of China (2019YFA0111800), Strategic Priority Research Programs of the Chinese Academy of Sciences (XDB29020000), National Natural Science Foundation of China (81922031, 82271790, 31770939), Key Research Program of Frontier Sciences of Chinese Academy of Sciences (ZDBS-LY-SM025), Fok Ying Tung Education Foundation to P.X., Youth Innovation Promotion Association of CAS to S.W..

## Author contributions

X.Q., F.Z. and W.L. designed and performed experiments and analyzed data; S.W. performed experiments and analyzed data; S.W. and P.X. initiated the study, designed and performed experiments, analyzed data, and wrote the paper.

---

**Supplementary Figure 1.**
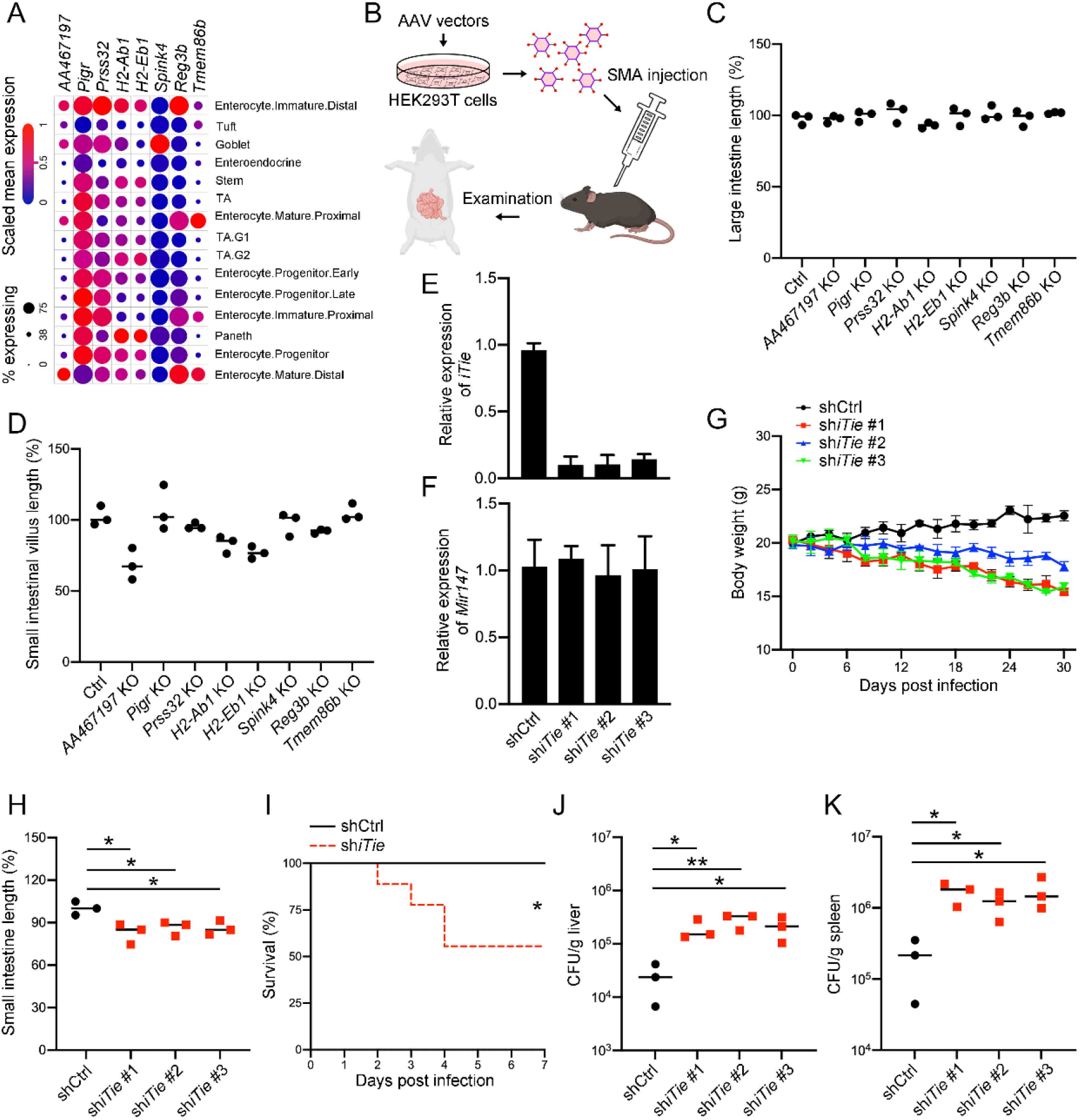
iTie is screened out as an essential player in the maintenance of intestinal tolerance. (A) Expression levels of the indicated genes in subsets of small intestinal epithelium were derived from the Single Cell Portal using the GSE92332 dataset. (B) A scheme was drawn for the AAV-based knockout or knockdown procedures. Briefly, adeno-associated virus (AAV) vectors containing gRNAs or shRNAs were packaged into viral particles in HEK293T cells, followed by recipient mice injection through the superior mesenteric artery. Mice were examined later. (C, D) One month post infection of gRNA-containing AAV viruses, mice were examined by quantifying the relative length of large intestines (C). Mouse small intestines were sectioned and stained with hematoxylin-eosin, followed by calculating the length changes of small intestinal villi (D). Three gRNAs were designed for each gene and one mouse was used for each gRNA. (E, F) shRNAs specific for *iTie* were packaged into AAV viral particles, followed by viral particle delivery into mice through the superior mesenteric artery. One month later, small intestinal enterocytes were collected and subjected to RP-PCR analyses using primers specific for *iTie* or *Mir147*. (G, H) shRNAs specific for *iTie* were packaged into AAV viral particles, followed by viral particle delivery into mice through the superior mesenteric artery. Body weights were monitored in the following month (G). Mouse small intestines were also sectioned and stained with hematoxylin-eosin, followed by calculating the length changes of small intestines (H). Three shRNAs were designed and three mice were used for each shRNA. (I–K) One month post viral infections of control or *iTie* specific shRNAs, mice were intragastrically gavaged with 4 x 10^9^ *L. monocytogenes*. Survival of the indicated mice were calculated in the following 7 days (I, n=9). Bacterial loads in livers (J) and spleens (K) were examined 3 days post bacterial infection. For E–K, three shRNAs were designed for *iTie* and three mice were used for each shRNA. Data were shown as means±SD. *, *P*<0.05; **, *P*<0.01. Experiments were repeated three times with similar results.

**Supplementary Figure 2.**
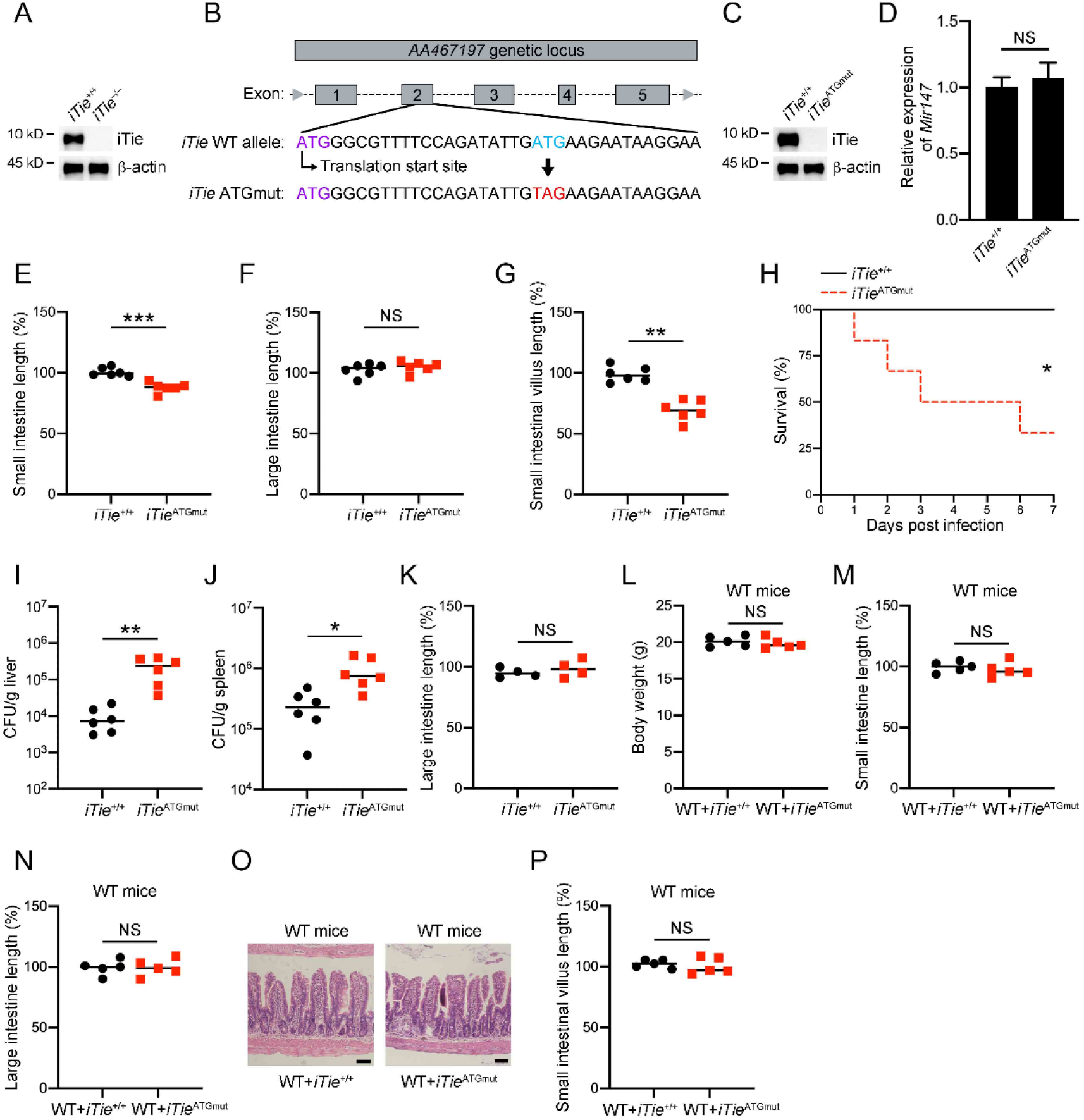
iTie is involved in the establishment of intestinal tolerance. (A) Small intestinal enterocytes were isolated from *iTie*^+/+^ and *iTie*^−/−^ mice and subjected to immunoblotting with antibodies against the indicated proteins. (B) A genetic manipulation strategy generating *iTie*^ATGmut^ mice were shown. Briefly, the 8^th^ genetic codon (ATG) of *iTie* was mutated to a stop codon (TAG) through the CRISPR/Cas9 technology. (C) Small intestinal enterocytes were isolated from *iTie*^+/+^ and *iTie*^ATGmut^ mice and subjected to immunoblotting with antibodies against the indicated proteins. (D) Enterocytes were isolated from *iTie*^+/+^ and *iTie*^ATGmut^ small intestines and subjected to RT-PCR analysis with *Mir147* specific primers. (E–G) The relative length changes of small intestines (E), large intestines (F) and small intestinal villi (G) between *iTie*^+/+^ and *iTie*^ATGmut^ mice were examined at 8 weeks of age. (H–J) *iTie*^+/+^ and *iTie*^ATGmut^ mice were intragastrically gavaged with 4 x 10^9^ *L. monocytogenes*. Survival of the indicated mice were calculated in the following 7 days (H, n=6). Bacterial loads in livers (I) and spleens (J) were examined 3 days post bacterial infection. (K) *iTie*^+/+^ and *iTie*^ATGmut^ mice were bred with antibiotics. Large intestines of *iTie*^+/+^ and *iTie*^ATGmut^ mice at 8 weeks of age were separated and the relative length changes were examined. (L–P) Newborn WT mice were bred in the same cages with *iTie*^+/+^ or *iTie*^ATGmut^ mice under SPF conditions. Body weights of WT mice were checked at 8 weeks of age (L). Small intestines (M) and large intestines (N) of the cohoused WT mice were separated and the relative length changes were calculated. Small intestines of the co-bred WT mice were also sectioned and stained with hematoxylin-eosin (O) and the relative length changes of small intestinal villi were calculated (P). Scale bar, 50 μm. NS, non-significant. Data were shown as means±SD. *, *P*<0.05; **, *P*<0.01; ***, *P*<0.001. Experiments were repeated three times with similar results.

**Supplementary Figure 3.**
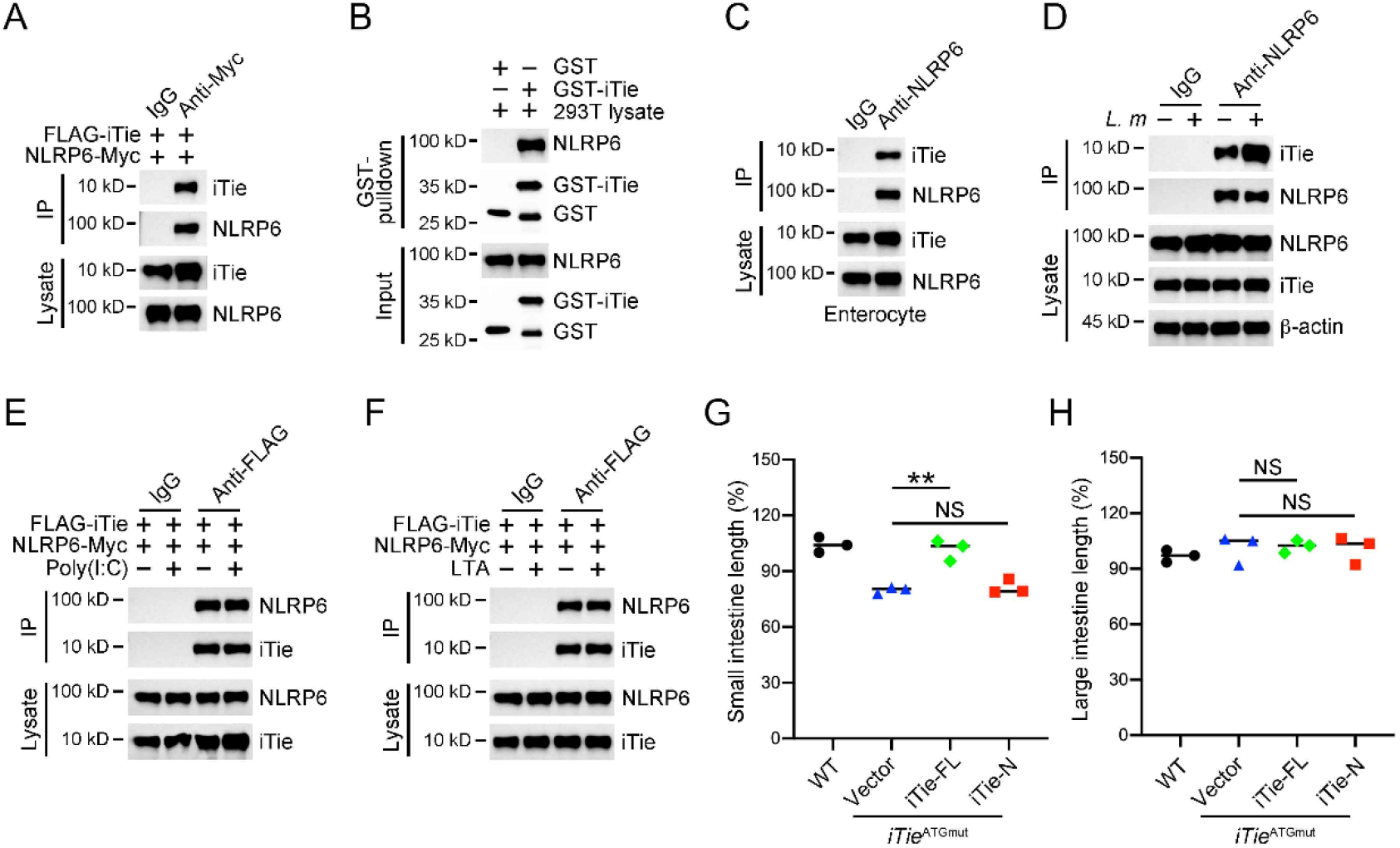
NLRP6 binds stronger with iTie than with NLRP6 ligands. (A) Plasmids encoding FLAG-tagged iTie and Myc-tagged NLRP6 were co-transfected into HEK293T cells for 24 h, followed by immunoprecipitation with antibody against Myc or with a control IgG. Immunoprecipitations and cell lysates were immunoblotted with antibodies against the indicated proteins. (B) Recombinant GST-iTie or control GST protein were incubated with HEK293T cell lysates containing overexpressed NLRP6 for 4 h, followed by a GST pull-down assay. Precipitates were immunoblotted with antibodies against the indicated proteins. (C) Enterocytes separated from the small intestine of WT mice were lysed and immunoprecipitated with antibody against NLRP6 or with a control IgG. Precipitates were immunoblotted with antibodies against the indicated proteins. (D) Enterocytes from the small intestine of WT mouse were incubated with *L. monocytogenes* at an MOI of 10 for 1.5 h. Cells were then supplemented with fresh medium containing 100 μg/ml Gentamycin and cultured for 16 h. Cells were then lysed and immunoprecipitated with antibody against NLRP6 or with a control IgG. Precipitates were immunoblotted with antibodies against the indicated proteins. (E, F) Plasmids encoding FLAG-tagged iTie and Myc-tagged NLRP6 were co-transfected into HEK293T cells for 24 h. Cells were further transfected with 1 μg/ml poly(I:C) (E) or 10 μg/ml LTA (F) for 4 h, followed by immunoprecipitation with antibody against FLAG or with a control IgG. Immunoprecipitations and cell lysates were immunoblotted with antibodies against the indicated proteins. (G, H) AAV viral particles containing gene fragments of iTie full-length or iTie N terminus (aa 1–41) were delivered into *iTie*^ATGmut^ mice through the superior mesenteric artery. One month later, the relative length changes of small intestines (G) and large intestines (H) were calculated. NS, non-significant. Data were shown as means±SD. **, *P*<0.01. Experiments were repeated three times with similar results.

**Supplementary Figure 4.**
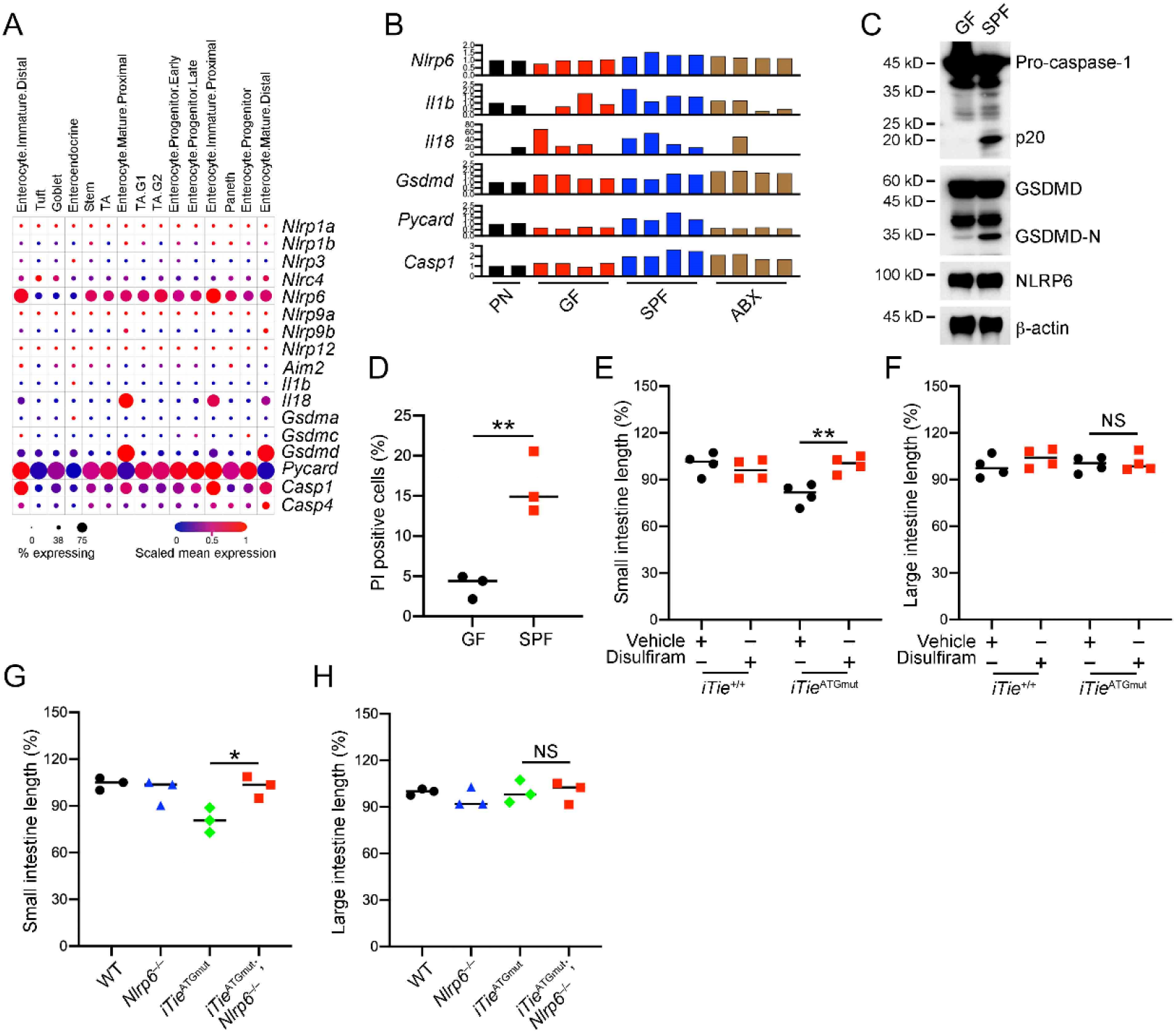
iTie inhibits NLRP6 inflammasome activation in enterocytes. (A) Expression levels of inflammasome-related genes in subsets of small intestinal epithelium were derived from the Single Cell Portal using the GSE92332 dataset. (B) Fold changes of expression levels of the indicated genes were calculated using RNA sequencing data of the indicated mouse intestine cells. The first sample of the PN group was set as 1. (C, D) Enterocytes were isolated from the small intestines of WT mice housed under germ-free or SPF conditions. Cells were either subjected to immunoblotting with antibodies against the indicated proteins (C) or stained with PI to calculate the percentages of PI positive cells (D). (E, F) *iTie*^+/+^ and *iTie*^ATGmut^ mice were administrated with 150 μg/ml disulfiram through drinking water for one month. The relative length changes of small intestines (E) and large intestines (F) were calculated. (G, H) The relative length changes of small intestines (G) and large intestines (H) were calculated among 8 weeks-old mice of WT, *Nlrp6*^−/−^, *iTie*^ATGmut^ and *iTie*^ATGmut^;*Nlrp6*^−/−^. NS, non-significant. Data were shown as means±SD. *, *P*<0.05; **, *P*<0.01; ***, *P*<0.001. Experiments were repeated three times with similar results.

**Supplementary Figure 5.**
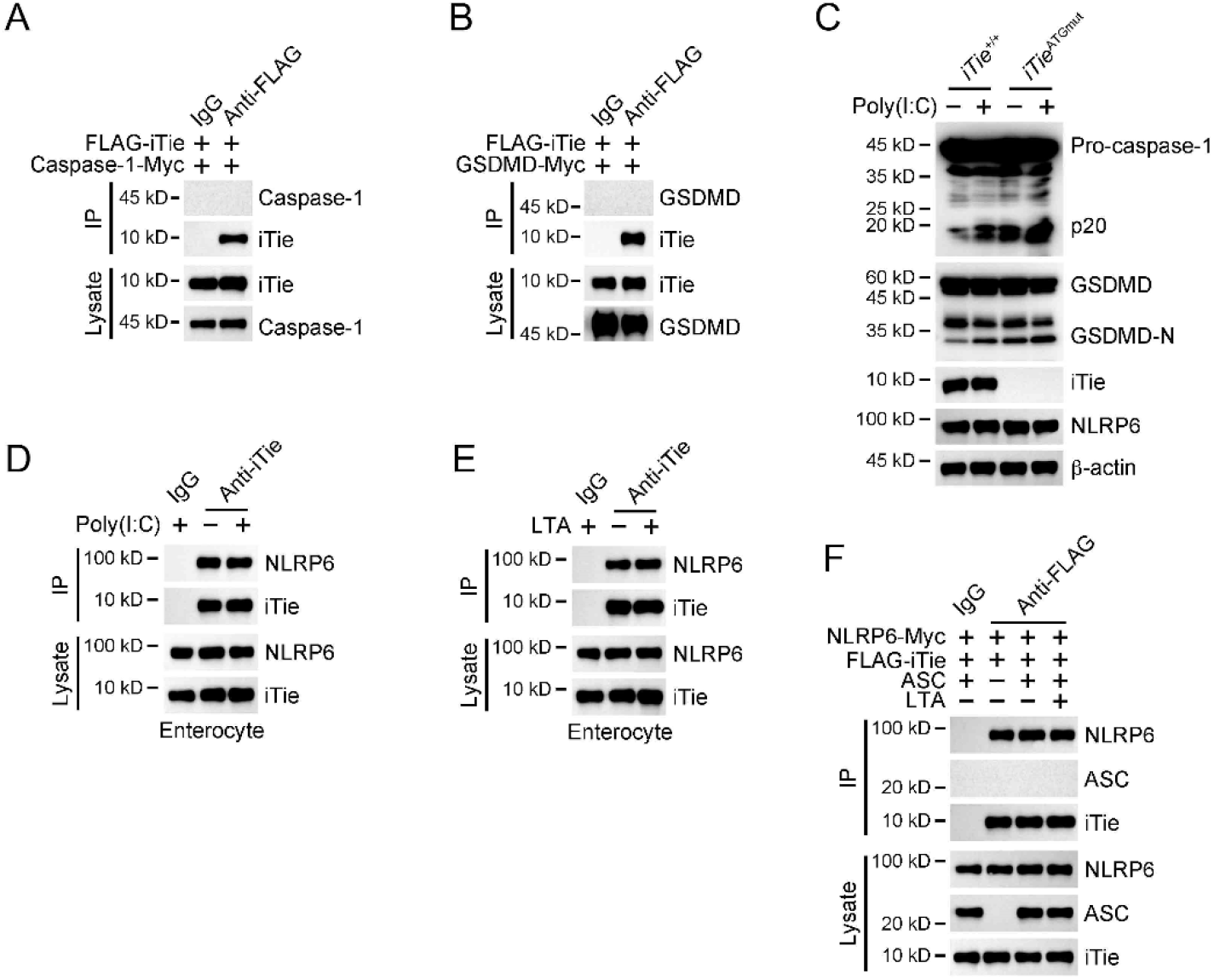
iTie prohibits the association between NLRP6 and ASC. (A) Plasmids encoding FLAG-tagged iTie and Myc-tagged caspase-1 were co-transfected into HEK293T cells for 24 h, followed by immunoprecipitation with antibody against FLAG or with a control IgG. Immunoprecipitations and cell lysates were immunoblotted with antibodies against the indicated proteins. (B) Plasmids encoding FLAG-tagged iTie and Myc-tagged GSDMD were co-transfected into HEK293T cells for 24 h, followed by immunoprecipitation with antibody against FLAG or with a control IgG. Immunoprecipitations and cell lysates were immunoblotted with antibodies against the indicated proteins. (C) Enterocytes were separated from the small intestines of *iTie*^+/+^ and *iTie*^ATGmut^ mice. Cells were transfected with 1 μg/ml poly(I:C) for 4 h, followed by immunoblotting with antibodies against the indicated proteins. (D) Enterocytes were separated from the small intestines of WT mice. Cells were transfected with 1 μg/ml poly(I:C) for 4 h, followed by immunoprecipitation with antibody against iTie or with a control IgG. Precipitates were immunoblotted with antibodies against the indicated proteins. (E) Enterocytes were separated from the small intestines of WT mice. Cells were transfected with 10 μg/ml LTA for 4 h, followed by immunoprecipitation with antibody against iTie or with a control IgG. Precipitates were immunoblotted with antibodies against the indicated proteins. (F) Plasmids encoding FLAG-tagged iTie, Myc-tagged NLRP6 and ASC were cotransfected into HEK293T cells for 24 h. Cells were then transfected with 10 μg/ml LTA for 4 h, followed by immunoprecipitation with antibody against FLAG or with a control IgG. Precipitates were immunoblotted with antibodies against the indicated proteins. Experiments were repeated three times with similar results.

**Supplementary Figure 6.**
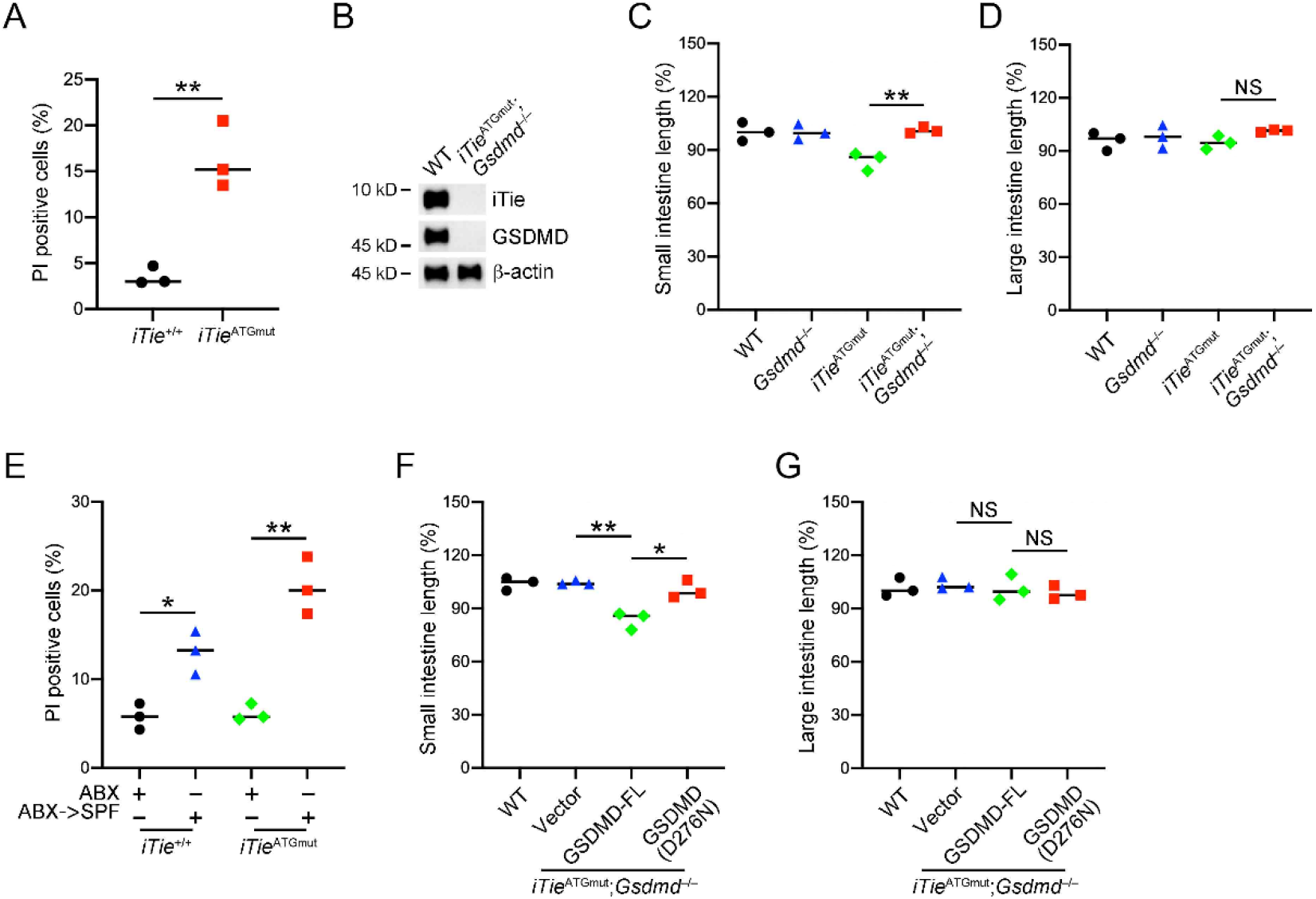
iTie maintains small intestinal tolerance through prohibiting GSDMD-mediated pyroptosis. (A) Enterocytes were separated from small intestines of *iTie*^+/+^ and *iTie*^ATGmut^ mice maintained in SPF conditions at 8 weeks of age. Cells were then stained with PI and the percentages of PI positive cells were calculated. (B) Enterocytes were separated from small intestines of WT and *iTie*^ATGmut^;*Gsdmd*^−/−^ mice at 8 weeks of age. Cells were lysed and immunoblotted with antibodies against the indicated proteins. (C, D) The relative length changes of small intestines (C) and large intestines (D) were calculated among 8 weeks-old mice of WT, *Gsdmd*^−/−^, *iTie*^ATGmut^ and *iTie*^ATGmut^;*Gsdmd*^−/−^. (E) 8 weeks-old *iTie*^+/+^ and *iTie*^ATGmut^ mice maintained in SPF conditions with antibiotics were transferred to SPF conditions without antibiotics for a month. Enterocytes were separated from the indicated mice and stained with PI to calculate the percentages of PI positive cells. (F, G) AAV viral particles containing gene fragments of WT GSDMD or mutant GSDMD (D276N) were delivered into *iTie*^ATGmut^;*Gsdmd*^−/−^ mice through the superior mesenteric artery. One month later, The relative length changes of small intestines (F) and large intestines (G) were examined. NS, non-significant. Data were shown as means±SD. *, *P*<0.05; **, *P*<0.01. Experiments were repeated three times with similar results.

## References

Allaire, J.M., Crowley, S.M., Law, H.T., Chang, S.Y., Ko, H.J., and Vallance, B.A. (2018). The Intestinal Epithelium: Central Coordinator of Mucosal Immunity. Trends Immunol 39, 677–696.

Birchenough, G.M., Nyström, E.E., Johansson, M.E., and Hansson, G.C. (2016). A sentinel goblet cell guards the colonic crypt by triggering Nlrp6-dependent Muc2 secretion. Science 352, 1535–1542.

Broz, P., Pelegrín, P., and Shao, F. (2020). The gasdermins, a protein family executing cell death and inflammation. Nat Rev Immunol 20, 143–157.

Bryce, P.J. (2016). Balancing Tolerance or Allergy to Food Proteins. Trends Immunol 37, 659–667.

Chen, G.Y., Liu, M., Wang, F., Bertin, J., and Núñez, G. (2011). A functional role for Nlrp6 in intestinal inflammation and tumorigenesis. J Immunol 186, 7187–7194.

Christgen, S., Place, D.E., and Kanneganti, T.D. (2020). Toward targeting inflammasomes: insights into their regulation and activation. Cell Res 30, 315–327.

Clayton, S.A., Daley, K.K., MacDonald, L., Fernandez-Vizarra, E., Bottegoni, G., O’Neil, J.D., Major, T., Griffin, D., Zhuang, Q., Adewoye, A.B., et al. (2021). Inflammation causes remodeling of mitochondrial cytochrome c oxidase mediated by the bifunctional gene C15orf48. Sci Adv 7, eabl5182.

Consortium, H.M.P. (2012). Structure, function and diversity of the healthy human microbiome. Nature 486, 207–214.

Ding, J., Wang, K., Liu, W., She, Y., Sun, Q., Shi, J., Sun, H., Wang, D.C., and Shao, F. (2016). Poreforming activity and structural autoinhibition of the gasdermin family. Nature 535, 111–116.

Elinav, E., Strowig, T., Kau, A.L., Henao-Mejia, J., Thaiss, C.A., Booth, C.J., Peaper, D.R., Bertin, J., Eisenbarth, S.C., Gordon, J.I., and Flavell, R.A. (2011). NLRP6 inflammasome regulates colonic microbial ecology and risk for colitis. Cell 145, 745–757.

Endou, M., Yoshida, K., Hirota, M., Nakajima, C., Sakaguchi, A., Komatsubara, N., and Kurihara, Y. (2020). Coxfa4l3, a novel mitochondrial electron transport chain Complex 4 subunit protein, switches from Coxfa4 during spermatogenesis. Mitochondrion 52, 1–7.

Evavold, C.L., and Kagan, J.C. (2019). Inflammasomes: Threat-Assessment Organelles of the Innate Immune System. Immunity 51, 609–624.

Fagerberg, L., Hallström, B.M., Oksvold, P., Kampf, C., Djureinovic, D., Odeberg, J., Habuka, M., Tahmasebpoor, S., Danielsson, A., Edlund, K., et al. (2014). Analysis of the human tissue-specific expression by genome-wide integration of transcriptomics and antibody-based proteomics. Mol Cell Proteomics 13, 397–406.

Floyd, B.J., Wilkerson, E.M., Veling, M.T., Minogue, C.E., Xia, C., Beebe, E.T., Wrobel, R.L., Cho, H., Kremer, L.S., Alston, C.L., et al. (2016). Mitochondrial Protein Interaction Mapping Identifies Regulators of Respiratory Chain Function. Mol Cell 63, 621–632.

Franchi, L., Kamada, N., Nakamura, Y., Burberry, A., Kuffa, P., Suzuki, S., Shaw, M.H., Kim, Y.G., and Núñez, G. (2012). NLRC4-driven production of IL-1β discriminates between pathogenic and commensal bacteria and promotes host intestinal defense. Nat Immunol 13, 449–456.

Glocker, E.O., Kotlarz, D., Boztug, K., Gertz, E.M., Schäffer, A.A., Noyan, F., Perro, M., Diestelhorst, J., Allroth, A., Murugan, D., et al. (2009). Inflammatory bowel disease and mutations affecting the interleukin-10 receptor. N Engl J Med 361, 2033–2045.

Guma, M., Stepniak, D., Shaked, H., Spehlmann, M.E., Shenouda, S., Cheroutre, H., Vicente-Suarez, I., Eckmann, L., Kagnoff, M.F., and Karin, M. (2011). Constitutive intestinal NF-κB does not trigger destructive inflammation unless accompanied by MAPK activation. J Exp Med 208, 1889–1900.

Hara, H., Seregin, S.S., Yang, D., Fukase, K., Chamaillard, M., Alnemri, E.S., Inohara, N., Chen, G.Y., and Núñez, G. (2018). The NLRP6 Inflammasome Recognizes Lipoteichoic Acid and Regulates Gram-Positive Pathogen Infection. Cell 175, 1651–1664.e1614.

Hooper, L.V., Littman, D.R., and Macpherson, A.J. (2012). Interactions between the microbiota and the immune system. Science 336, 1268–1273.

Hu, J.J., Liu, X., Xia, S., Zhang, Z., Zhang, Y., Zhao, J., Ruan, J., Luo, X., Lou, X., Bai, Y., et al. (2020). FDA-approved disulfiram inhibits pyroptosis by blocking gasdermin D pore formation. Nat Immunol 21, 736–745.

Iweala, O.I., and Nagler, C.R. (2019). The Microbiome and Food Allergy. Annu Rev Immunol 37, 377–403.

Johansson, M.E., Larsson, J.M., and Hansson, G.C. (2011). The two mucus layers of colon are organized by the MUC2 mucin, whereas the outer layer is a legislator of host-microbial interactions. Proc Natl Acad Sci U S A 108 Suppl 1, 4659–4665.

Kayama, H., Okumura, R., and Takeda, K. (2020). Interaction Between the Microbiota, Epithelia, and Immune Cells in the Intestine. Annu Rev Immunol 38, 23–48.

Knodler, L.A., Crowley, S.M., Sham, H.P., Yang, H., Wrande, M., Ma, C., Ernst, R.K., Steele-Mortimer, O., Celli, J., and Vallance, B.A. (2014). Noncanonical inflammasome activation of caspase-4/caspase-11 mediates epithelial defenses against enteric bacterial pathogens. Cell Host Microbe 16, 249–256.

Lamkanfi, M., and Dixit, V.M. (2014). Mechanisms and functions of inflammasomes. Cell 157, 1013–1022.

Lee, C.Q.E., Kerouanton, B., Chothani, S., Zhang, S., Chen, Y., Mantri, C.K., Hock, D.H., Lim, R., Nadkarni, R., Huynh, V.T., et al. (2021). Coding and non-coding roles of MOCCI (C15ORF48) coordinate to regulate host inflammation and immunity. Nat Commun 12, 2130.

Levy, M., Shapiro, H., Thaiss, C.A., and Elinav, E. (2017). NLRP6: A Multifaceted Innate Immune Sensor. Trends Immunol 38, 248–260.

Levy, M., Thaiss, C.A., Zeevi, D., Dohnalová, L., Zilberman-Schapira, G., Mahdi, J.A., David, E., Savidor, A., Korem, T., Herzig, Y., et al. (2015). Microbiota-Modulated Metabolites Shape the Intestinal Microenvironment by Regulating NLRP6 Inflammasome Signaling. Cell 163, 1428–1443.

Li, R., and Zhu, S. (2020). NLRP6 inflammasome. Mol Aspects Med 76, 100859.

Lipinski, S., Grabe, N., Jacobs, G., Billmann-Born, S., Till, A., Häsler, R., Aden, K., Paulsen, M., Arlt, A., Kraemer, L., et al. (2012). RNAi screening identifies mediators of NOD2 signaling: implications for spatial specificity of MDP recognition. Proc Natl Acad Sci U S A 109, 21426–21431.

Liu, X., Zhang, Z., Ruan, J., Pan, Y., Magupalli, V.G., Wu, H., and Lieberman, J. (2016). Inflammasome-activated gasdermin D causes pyroptosis by forming membrane pores. Nature 535, 153–158.

Martens, E.C., Neumann, M., and Desai, M.S. (2018). Interactions of commensal and pathogenic microorganisms with the intestinal mucosal barrier. Nat Rev Microbiol 16, 457–470.

Mowat, A.M. (2018). To respond or not to respond - a personal perspective of intestinal tolerance. Nat Rev Immunol 18, 405–415.

Mukherjee, S., Kumar, R., Tsakem Lenou, E., Basrur, V., Kontoyiannis, D.L., Ioakeimidis, F., Mosialos, G., Theiss, A.L., Flavell, R.A., and Venuprasad, K. (2020). Deubiquitination of NLRP6 inflammasome by Cyld critically regulates intestinal inflammation. Nat Immunol 21, 626–635.

Okumura, R., and Takeda, K. (2017). Roles of intestinal epithelial cells in the maintenance of gut homeostasis. Exp Mol Med 49, e338.

Patankar, J.V., and Becker, C. (2020). Cell death in the gut epithelium and implications for chronic inflammation. Nat Rev Gastroenterol Hepatol 17, 543–556.

Perez-Lopez, A., Behnsen, J., Nuccio, S.P., and Raffatellu, M. (2016). Mucosal immunity to pathogenic intestinal bacteria. Nat Rev Immunol 16, 135–148.

Peterson, L.W., and Artis, D. (2014). Intestinal epithelial cells: regulators of barrier function and immune homeostasis. Nat Rev Immunol 14, 141–153.

Polyak, S., Mah, C., Porvasnik, S., Herlihy, J.D., Campbell-Thompson, M., Byrne, B.J., and Valentine, J.F. (2008). Gene delivery to intestinal epithelial cells in vitro and in vivo with recombinant adeno-associated virus types 1, 2 and 5. Dig Dis Sci 53, 1261–1270.

Rathinam, V.A., and Fitzgerald, K.A. (2016). Inflammasome Complexes: Emerging Mechanisms and Effector Functions. Cell 165, 792–800.

Sellin, M.E., Müller, A.A., Felmy, B., Dolowschiak, T., Diard, M., Tardivel, A., Maslowski, K.M., and Hardt, W.D. (2014). Epithelium-intrinsic NAIP/NLRC4 inflammasome drives infected enterocyte expulsion to restrict Salmonella replication in the intestinal mucosa. Cell Host Microbe 16, 237–248.

Sender, R., Fuchs, S., and Milo, R. (2016). Are We Really Vastly Outnumbered? Revisiting the Ratio of Bacterial to Host Cells in Humans. Cell 164, 337–340.

Shen, C., Li, R., Negro, R., Cheng, J., Vora, S.M., Fu, T.M., Wang, A., He, K., Andreeva, L., Gao, P., et al. (2021). Phase separation drives RNA virus-induced activation of the NLRP6 inflammasome. Cell 184, 5759–5774.e5720.

Shi, J., Zhao, Y., Wang, K., Shi, X., Wang, Y., Huang, H., Zhuang, Y., Cai, T., Wang, F., and Shao, F. (2015). Cleavage of GSDMD by inflammatory caspases determines pyroptotic cell death. Nature 526, 660–665.

Sun, Y., Zhang, M., Chen, C.C., Gillilland, M., 3rd, Sun, X., El-Zaatari, M., Huffnagle, G.B., Young, V.B., Zhang, J., Hong, S.C., et al. (2013). Stress-induced corticotropin-releasing hormone-mediated NLRP6 inflammasome inhibition and transmissible enteritis in mice. Gastroenterology 144, 1478–1487, 1487.e1471-1478.

Wang, K., Sun, Q., Zhong, X., Zeng, M., Zeng, H., Shi, X., Li, Z., Wang, Y., Zhao, Q., Shao, F., and Ding, J. (2020). Structural Mechanism for GSDMD Targeting by Autoprocessed Caspases in Pyroptosis. Cell 180, 941–955.e920.

Wang, P., Zhu, S., Yang, L., Cui, S., Pan, W., Jackson, R., Zheng, Y., Rongvaux, A., Sun, Q., Yang, G., et al. (2015). Nlrp6 regulates intestinal antiviral innate immunity. Science 350, 826–830.

Wlodarska, M., Thaiss, C.A., Nowarski, R., Henao-Mejia, J., Zhang, J.P., Brown, E.M., Frankel, G., Levy, M., Katz, M.N., Philbrick, W.M., et al. (2014). NLRP6 inflammasome orchestrates the colonic host-microbial interface by regulating goblet cell mucus secretion. Cell 156, 1045–1059.

Xiao, H., Gulen, M.F., Qin, J., Yao, J., Bulek, K., Kish, D., Altuntas, C.Z., Wald, D., Ma, C., Zhou, H., et al. (2007). The Toll-interleukin-1 receptor member SIGIRR regulates colonic epithelial homeostasis, inflammation, and tumorigenesis. Immunity 26, 461–475.

Zhou, J., Wang, H., Lu, A., Hu, G., Luo, A., Ding, F., Zhang, J., Wang, X., Wu, M., and Liu, Z. (2002). A novel gene, NMES1, downregulated in human esophageal squamous cell carcinoma. Int J Cancer 101, 311–316.

